# Solid-State NMR Analysis of *Schizosaccharomyces pombe* Reveals Role of *α*-Amylase Family Enzymes in Cell Wall Structure and Function

**DOI:** 10.1101/2025.03.10.642465

**Authors:** Anand Jacob, Alaina H. Willet, Maya G. Igarashi, Mustapha El Hariri El Nokab, Lesley A. Turner, Abdulrahman Khalid A. Alsanad, Tuo Wang, Kathleen L. Gould

**Affiliations:** Department of Chemistry, Michigan State University, East Lansing, MI 48824, USA; Department of Cell and Developmental Biology, Vanderbilt University School of Medicine, Nashville, TN 37232

**Keywords:** *Schizosaccharomyces pombe*, solid-state NMR, cell division, amylase, cell wall, α- glucan

## Abstract

The fission yeast *Schizosaccharomyces pombe* is a widely employed model organism for studying the eukaryotic cell cycle. Like plants and bacteria, *S. pombe* must build a cell wall in concert with its cell cycle, but how cell wall-synthesizing and remodeling enzymes mediate this process remains unclear. Here we characterize the functions of Aah1 and Aah3, two related *S. pombe* α-amylases that are putative members of this evolutionarily conserved family of cell wall-modifying proteins. We found that unlike rod-shaped wildtype *S. pombe* cells, *aah1*Δ *aah3*Δ cells are nearly spherical, grow slowly, have thickened cell walls, and have severe defects in cell separation following cytokinesis. Solid-state NMR spectroscopy analyses of intact wildtype and *aah1*Δ *aah3*Δ cells revealed that *aah1*Δ *aah3*Δ cell walls are rigidified with a significant reduction in the α-glucan matrix, characterized by reduced amounts of the major α-1,3-glucan and the minor α-1,4-glucan within the rigid and mobile phases; this reduction was compensated for by a two-fold increase in β-glucan content. Indeed, viability of *aah1*Δ *aah3*Δ cells depended on β-glucan upregulation and the cell wall integrity pathway that mediates it. While *aah1*Δ *aah3*Δ cells resemble cells with impaired function of the transglycosylation domain of α-glucan synthase 1 (Ags1), increased expression of Aah3 does not compensate for impaired Ags1 function or vice-versa. Overall, our data suggest that Aah1 and Aah3 are required in addition to Ags1, likely downstream, for the transglycosylation of α-glucan chains to generate fibers of appropriate dimensions to support proper cell morphology, growth, and division.

**Significance Statement:** This study utilized a range of imaging techniques and high-resolution solid-state NMR spectroscopy of intact *S. pombe* cells to refine our understanding of *S. pombe* cell wall composition. This study also determined that two related GPI-anchored α-amylase family proteins, Aah1 and Aah3, likely act as transglycosylases non-redundantly with an α-glucan synthase in the synthesis of α-glucan chains of appropriate content and size to support polarized growth and cell division. Our results also highlight the anti-fungal therapeutic potential of GPI-anchored enzymes acting in concert with glucan synthases.

## INTRODUCTION

*Schizosaccharomyces pombe* is a rod-shaped unicellular eukaryote that grows by tip extension and divides medially using an actomyosin-based contractile apparatus and a septum (1–4). Its relatively simple genome and the conservation of key cellular processes between *S. pombe* and multicellular eukaryotes has made it a valuable model organism for providing critical insights into fundamental biological mechanisms such as cell cycle regulation applicable to higher organisms (5–7). *S. pombe* cells are encompassed by a cell wall that serves to counteract their high internal turgor pressure and protects them from environmental stresses (8). However, as for all cell-walled organisms, how cell wall synthesis and remodeling is regulated to simultaneously provide structural support and allow for cell growth and division is not fully understood (9–12).

The *S. pombe* cell wall is organized in three distinct layers, as visualized by electron microscopy. The layer most distal from the plasma membrane consists of galactomannan, the medial layer is mostly composed of α-1,3-glucan, β-1,3-glucan and β-1,6-glucan (**Fig. 1**), and the layer most proximal to the plasma membrane contains galactomannan and a dense meshwork of glucans (13–18). The galactomannan in *S. pombe* primarily consists of α-1,6-linked mannan backbones with branches formed by α-1,2 or α-1,3-mannose residues with a terminal galactose residue located on the non-reducing end (19–24). Galactomannans are synthesized by multiple non-essential proteins and cells lacking all of them remain viable (25). In contrast, most glucan synthases are essential for viability during vegetative growth. These include the α-glucan synthase Ags1, and the β-glucan synthases Bgs1, Bgs3, and Bgs4 (26–30).

**Figure 1.**
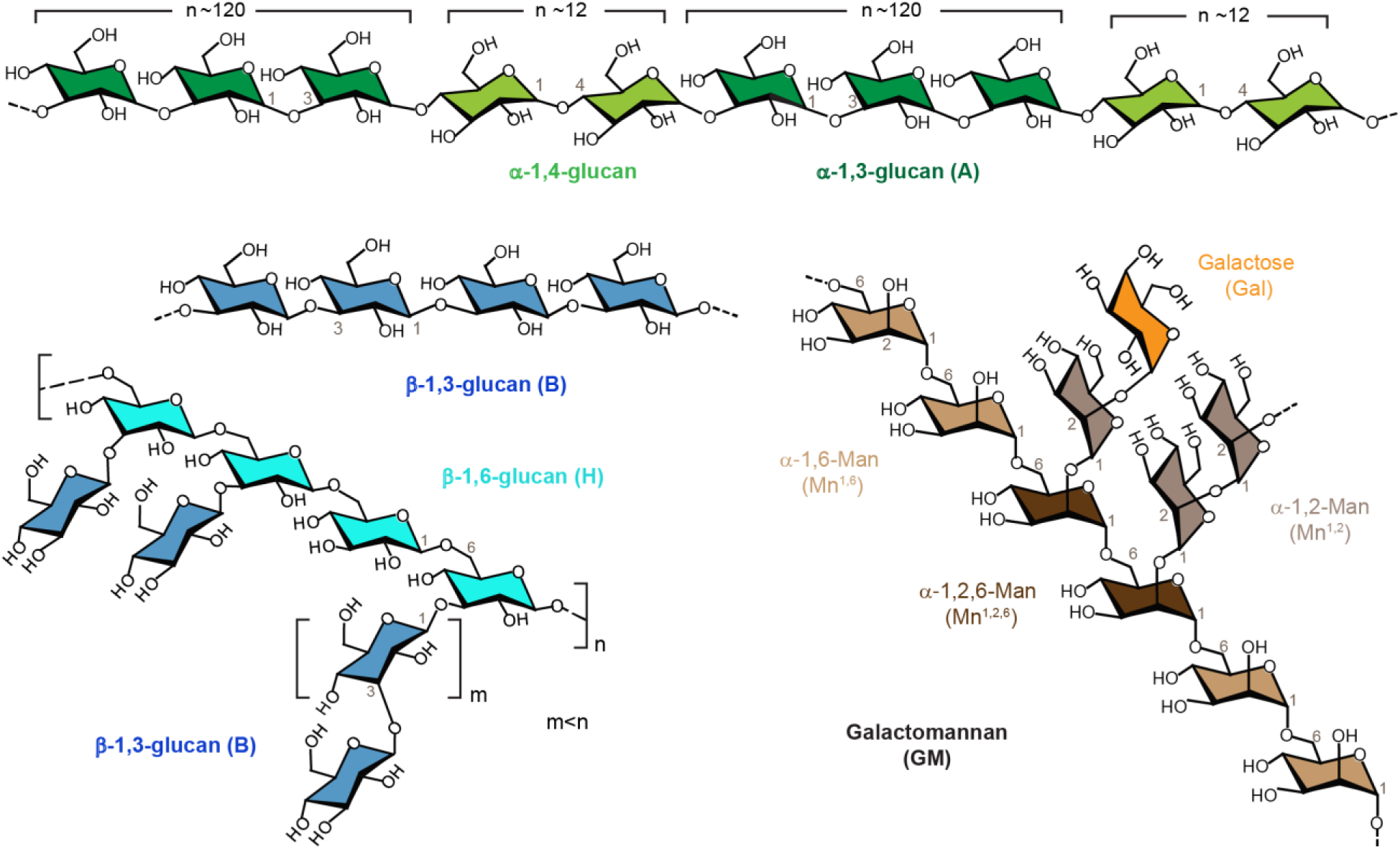
Carbohydrate structures in the cell wall of *S. pombe*. Simplified structural representations are provided for α-glucans (top), β-glucans (bottom left), and galactomannan (bottom right). Key linkage sites and NMR abbreviations are labelled.

Ags1 is a 12-pass transmembrane protein that contains an intracellular glycosyl transferase 5 (GT5) domain and an extracellular glycoside hydrolase 13 (GH13) domain (31). Both domains are essential for Ags1 function (30, 32). It is proposed that the GT5 domain synthesizes a linear ∼120 residue α-1,3-glucan chain from an α-1,4-glucan primer and these chains are extruded through the transmembrane channel (14, 30). Subsequently, the Ags1 GH13 domain is suggested to act as a transglycosylase that joins the ∼120 residue ɑ-glucan chains together to construct a ∼260 α-glucan chain (**Fig. 1**) (14). Cells defective in *ags1* GH13 domain function become rounded, have an abnormal and thickened cell wall, and eventually lyse and die (32–34).

GH13 domains are also found in α-amylases, an evolutionarily conserved protein family of which five of the seven in *S. pombe* (Aah1-4 and Mde5) are also glycosylphosphatidylinositol (GPI)- anchored cell-wall associated proteins (35–38). The GPI anchor presumably positions these proteins near their substrates on the outside of the plasma membrane (36, 39). One of the α- amylase-related proteins, Aah3, has previously been reported to be involved in maintaining the normal rod shape of *S. pombe* during vegetative growth (38). While α-amylases typically hydrolyze α-1,4-glucans, lysates of *S. pombe* cells and their cultured media lack detectable α-1,4- glucan glycosidase activity (40). Therefore, it has been suggested that like the GH13 domain within Ags1, Aah1-4 and Mde5 are not hydrolases but rather transglycosylases, another common catalytic function of GH13 domain containing enzymes (41, 42).

Here, we found that cells lacking the related Aah1 and Aah3 enzymes have drastic defects in cell shape; they are spherical rather than rod-shaped and fail to physically separate following cytokinesis, resulting in clumps of cells. Transmission electron microscopy (TEM) showed that *aah1*Δ *aah3*Δ cells have abnormally thick cell walls lacking the typical three-layered organization of wildtype cell walls. This phenotype is correlated with limited water permeability and significantly reduced cell wall dynamics on both micro- and nano-second timescales, as shown by solid-state NMR spectroscopy (43). Determination of cell wall composition by solid-state NMR uncovered previously unrecognized polymorphic forms of polysaccharides within the *S. pombe* cell wall and revealed that *aah1*Δ *aah3*Δ cells have a significant reduction in α-1,3-glucan and galactomannan content. There is also a compensatory increase in β-1,3-glucan content in comparison to wildtype cells due to activation of the cell wall integrity pathway. The phenotypes of *aah1*Δ *aah3*Δ cells are strikingly similar to cells with defective Ags1 GH13 domain function (33). However, overexpression of *aah3* failed to rescue defective *ags1* GH13 function and overexpression of *ags1* failed to rescue *aah1*Δ *aah3*Δ cell phenotypes. Thus, we propose that Aah1 and Aah3 are required in addition to Ags1 for transglycosylating α-glucan chains to create the appropriately sized fibers needed to maintain proper cell shape and allow for cell separation following cytokinesis.

## RESULTS

### Aah1 and Aah3 are important for proper cell shape and cell separation in *S. pombe*

To investigate the cellular functions of the putative GPI-anchored α-amylase proteins in *S. pombe,* we analyzed the growth of single and double gene deletion strains. We excluded *mde5* from our analysis due to its meiotic-specific expression (44). The *aah1Δ, aah2Δ,* and *aah4*Δ cells grew like wildtype at all temperatures tested, and *aah3*Δ cells were cold-sensitive (**Fig. S1**). Of the double mutant strains, only *aah1*Δ *aah3*Δ cells showed an enhanced growth defect at all temperatures tested (**Fig. S1**). Combining *aah2*Δ or *aah4*Δ with *aah1*Δ *aah3*Δ did not result in any further growth defects (**Fig. S1**). Thus, we concluded that Aah1 and Aah3 act redundantly and are the two *S. pombe* α-amylase homologs important for vegetative cell growth.

To better understand the influence of Aah1 and Aah3 on cell growth, we imaged wildtype, *aah1Δ, aah3Δ,* and *aah1*Δ *aah3*Δ cells and quantified the ratio of cell length to cell width at septation as a measurement of cell morphology. Wildtype and *aah1*Δ cells had an average length-to-width ratio of ∼4.5 indicative of their rod shape (**Fig. 2A, B**). *aah3*Δ cells were abnormally pear-shaped and had a lower length-to-width ratio (**Fig. 2A, B**) consistent with published work (38, 45). Lastly, *aah1*Δ *aah3*Δ cells were spherical rather than rod-shaped resulting in a ratio of septated cell length-to-width nearing one (**Fig. 2A, B**). Almost every *aah1*Δ *aah3*Δ cell also contained at least one septum, and the cells tended to form clumps (**Fig. 2A**). These observations were confirmed by fixing and staining the cells with DAPI and methyl blue to mark the DNA and the cell wall, respectively, and quantifying the number of nucleated cell compartments per clump or cell (**Fig. 2C, D**). While approximately 15% of wildtype, *aah1*Δ, and *aah3*Δ cells had two nucleated compartments, many *aah1*Δ *aah3*Δ clumps had three, four, or five nucleated compartments (**Fig. 2C, D**), consistent with a defect in cell separation following cytokinesis.

**Figure 2.**
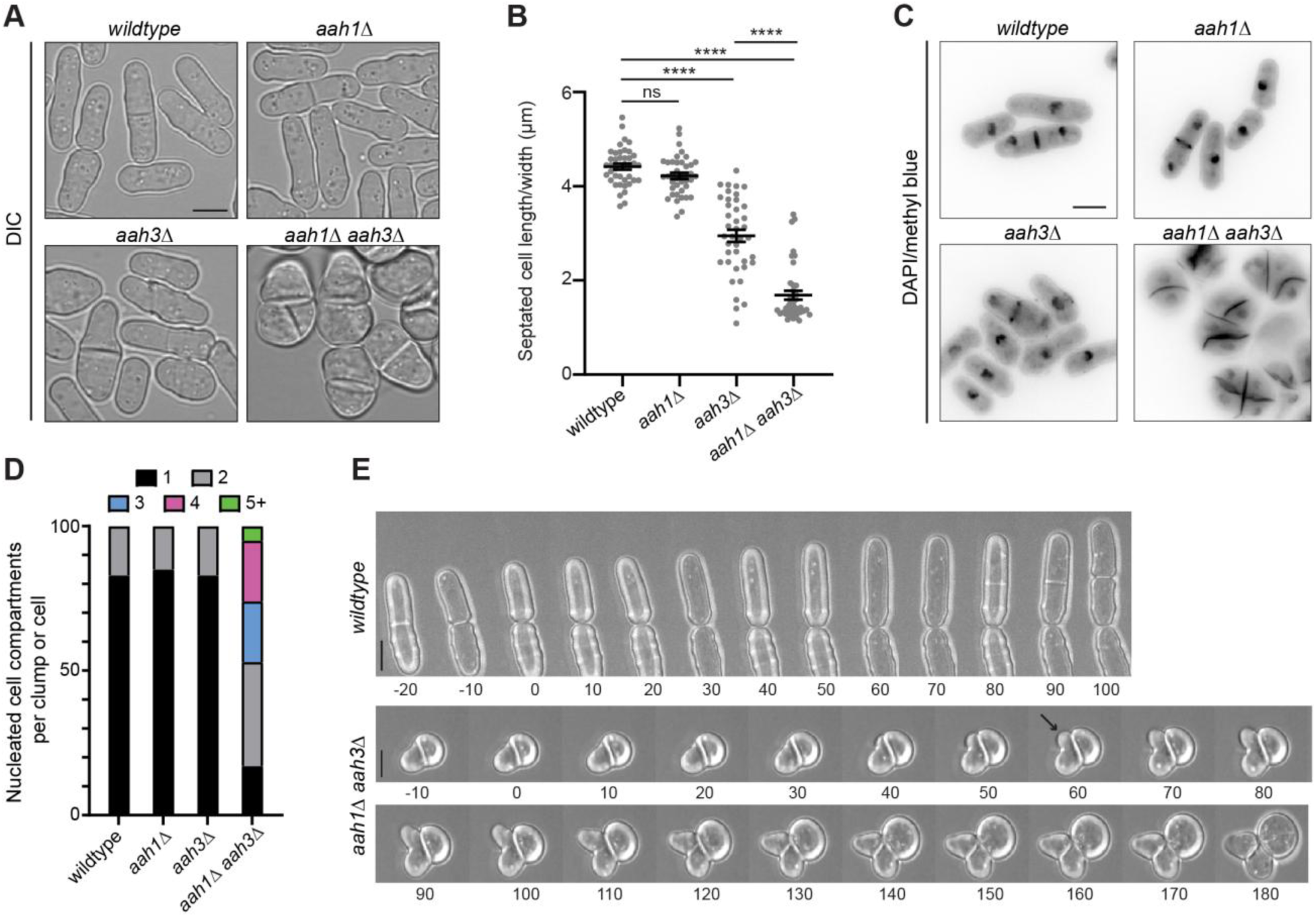
Aah1 and Aah3 are important for proper cell shape in *S. pombe*. (**A**) Representative images of the indicated strains grown at 32°C and imaged by differential interference contrast (DIC) microscopy. (**B**) Quantification of septated cell width divided by cell length. n=40 cells for each strain from two biological replicates. Horizontal bars represent the mean, and error bars represent the standard error. ****P≤0.0001 (one-way ANOVA with Tukey’s post-test). Wildtype versus *aah1*Δ, P = 0.436. (**C**) Representative images of the indicated strains grown at 32°C, fixed, and stained with DAPI and methyl blue. (**D**) Quantification of the number of nucleated compartments per clump. n=166 for *wildtype*, n=211 for *aah1*Δ, n=347 for *aah3*Δ and n=84 for *aah1*Δ *aah3*Δ from two independent experiments. (**E**) Live-cell time-lapse imaging of the indicated cells imaged by DIC every 5 min. Every other image is shown in the representative montages. Time zero in wildtype cells is the first frame of cell separation. Time zero is arbitrary for *aah1*Δ *aah3*Δ cells because they typically do not separate following cell division. The black arrow on the *aah1*Δ *aah3*Δ cell montage indicates a cell that initially polarizes at one cell end. Scale bars represent 5 μm for all panels.

To better understand the cell separation defect observed in *aah1*Δ *aah3*Δ cells, we performed differential interference contrast (DIC) live-cell time-lapse imaging. Wildtype cells septated and subsequently physically separated, as expected (**Fig. 2E**). In contrast, *aah1*Δ *aah3*Δ cells did not physically separate following what appeared to be the completion of septation (**Fig. 2E**). Many *aah1*Δ *aah3*Δ cells septated again while still being physically connected to their previous sister cell (**Fig. 2E**). Interestingly, some *aah1*Δ *aah3*Δ cells exhibited polarized growth initially after septation, indicating the cells maintained some polarity cues (**Fig. 2E**).

### *aah1*Δ *aah3*Δ cells show abnormal cell wall ultrastructure

To examine the ultrastructure of the cell wall, we analyzed wildtype and *aah1*Δ *aah3*Δ cells by transmission electron microscopy (TEM). Wildtype cells showed the characteristic uniform tri-layered cell wall structure that includes a more electron dense plasma membrane proximal galactomannan and glucan layer, a less dense medial glucan layer and an outermost dense galactomannan layer (**Fig. 3A, B**) (16). The cell wall of *aah1*Δ *aah3*Δ cells similarly included a plasma membrane proximal dense layer and a medial less dense layer however, many cells appeared to lack the outermost layer (**Fig. 3A, B**). Additional stratification could also be observed in some *aah1*Δ *aah3*Δ cells. Specifically, a more electron dense layer within the medial layer, referred to as an internal cell wall (ICW) remnant, was detected (**Fig. 3B**) (46). We further probed the observation that many *aah1*Δ *aah3*Δ cells lacked an outermost layer, typically comprised of galactomannans, by staining cells with the Concanavalin A lectin labeled with fluorescein isothiocyanate (FITC). This dye binds mannose residues on the cell surface and is a readout for the presence of the outermost galactomannan layer (47). Indeed, *aah1*Δ *aah3*Δ cells had reduced FITC-Concanavalin A staining compared to wildtype cells, aligning with the TEM observations (**Fig. S2**).

**Figure 3.**
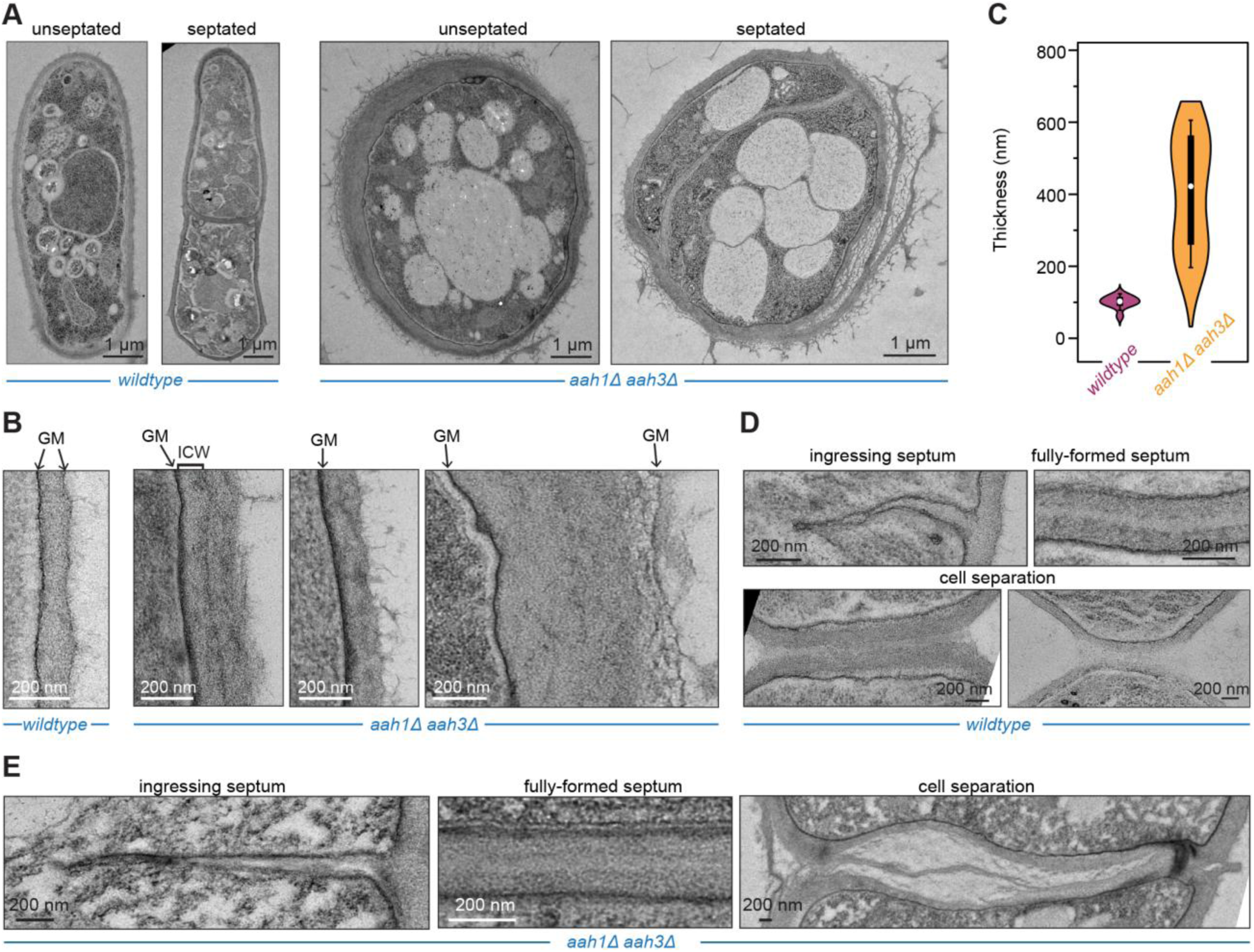
*aah1*Δ *aah3*Δ cells have thicker and abnormally structured cell walls. Representative transmission electron micrographs of wildtype and *aah1*Δ *aah3*Δ cells that were grown at 32°C in YE prior to freezing the cells live. (**A**) Examples of unseptated and septated cells from wildtype and *aah1*Δ *aah3*Δ cells with scale bars of 1 µm. (**B**) Zoomed in views of the lateral cell wall for each strain. Scale bars = 200 nm. Arrows indicate galactomannan (GM) cell wall layers. A black bracket indicates the location of the internal cell wall (ICW) in the *aah1*Δ *aah3*Δ mutant. (**C**) Violin plot of cell wall thickness measured using TEM images. Black rectangle denotes the interquartile range (IQR) from 25^th^ to 75^th^ percentile. n=100 for both wildtype and mutant cells. The white circle signifies the median of the dataset, while the black vertical line indicates the standard range of 1.5 IQR. (**D)** Wildtype and (**E**) *aah1*Δ *aah3*Δ mutant zoomed in views of ingressing septa, fully formed septa and septa in the process of degradation from each strain. Scale bars = 200 nm.

Despite the apparent lack of an outer galactomannan layer, the width of *aah1*Δ *aah3*Δ cell walls was more variable compared to wildtype cell walls and generally the walls were thicker (**Fig. 3A- C**), averaging approximately 400 nm in comparison to the average of approximately 100 nm for wildtype cell wall thickness (**Fig. 3C**). We also observed that the outermost layers of *aah1*Δ *aah3*Δ cell walls appeared to be abnormally shedding from the cell, a phenotype not observed in wildtype cells and possibly explaining the reduction of FITC-Concanavalin A staining (**Fig. 3A**).

For *S. pombe* cells to divide, an actin- and myosin-based cytokinetic ring is built at the cell equator that eventually constricts (48). Cell wall material is assembled behind the constricting cytokinetic ring to form both a primary septum that is eventually degraded and a secondary septum that becomes the new tips’ cell walls (49, 50). The primary septum is mainly composed of β-1,3- glucans while the secondary septum contains ɑ-1,3-glucans, branched and linear β-1,3-glucans, and galactomannans (17). Cell separation defects, like those of *aah1*Δ *aah3*Δ cells, can arise from either incomplete cytokinesis or incomplete primary septum degradation. To differentiate between these possibilities, we used TEM to visualize the septa of wildtype and *aah1*Δ *aah3*Δ cells. Septa that had not finished ingressing appeared similar between the samples (**Fig. 3D, E**). Further, fully formed septa showed a typical trilamellar structure in both cell types (**Fig 3A, D-E**). Thus, septum formation did not appear to be aberrant in *aah1*Δ *aah3*Δ cells. However, differences between wildtype and *aah1*Δ *aah3*Δ division sites were visualized in separating daughter cells. Wildtype daughter cells began separation from the lateral cell sides. In contrast, *aah1*Δ *aah3*Δ daughter cells maintained connections at their lateral cell edges while gaps developed where the primary septum had been (**Fig. 3D, E**). This suggests that while at least the majority of the primary septum had been degraded, abnormal connections at the lateral sides prevented cell separation.

### *aah1*Δ *aah3*Δ cells have altered cell wall composition and rigidity

Because Aah1 and Aah3 are predicted cell wall modifying enzymes, we used solid-state NMR to compare the cell wall composition and polysaccharide structure between wildtype and *aah1*Δ *aah3*Δ cells. This spectroscopic approach has not previously been applied to intact fission yeast and offers several technical advantages, including the ability to directly analyze cell walls in intact, natively hydrated cells (43, 51–53). Additionally, it provides spatial information on cell wall composition across different domains with distinct dynamics and enables the detection of polymorphic polysaccharide forms (54). Polymorphs, which are chemically identical but structurally distinct, commonly occur in cellular environments and can arise due to variations in conformation, hydrogen-bonding patterns, or ligand binding (54–59).

Initial 1D solid-state NMR screening of intact wildtype and *aah1*Δ *aah3*Δ cells revealed significant alterations in the polysaccharide content and the polysaccharide distribution across dynamically distinct domains of the *S. pombe* cell wall (**Fig. 4A**). The rigid cell wall phase is typically attributed to molecules packed with chitin microfibrils in other fungal species. In *S. pombe*, however, this phase is likely formed by tightly clustered molecules and is predominantly composed of β-1,3- glucan and α-1,3-glucan, with evidence of polymorphic forms of α-1,3-glucan present in the wildtype, exemplified by two carbon 1 peaks at 101 and 100 ppm (A^a^1 and A^b^1) (**Fig. 4A, B**). In comparison, the *aah1*Δ *aah3*Δ mutant exhibited a significantly reduced α-1,3-glucan content, with the A^a^1 peak diminished to one-third of its original intensity and the A^b^1 peak completely absent (**Fig. 4C**). These α-1,3-glucans and β-1,3-glucans were also detected in the mobile phase originating from the surface and membrane-proximal layers, which are rich in highly dynamic galactomannan and glycoproteins. The mobile phase additionally contained contributions from β- 1,6-glucan and α-1,2- and α-1,6-linked mannose units in mannan (**Fig. 4A**).

**Figure 4.**
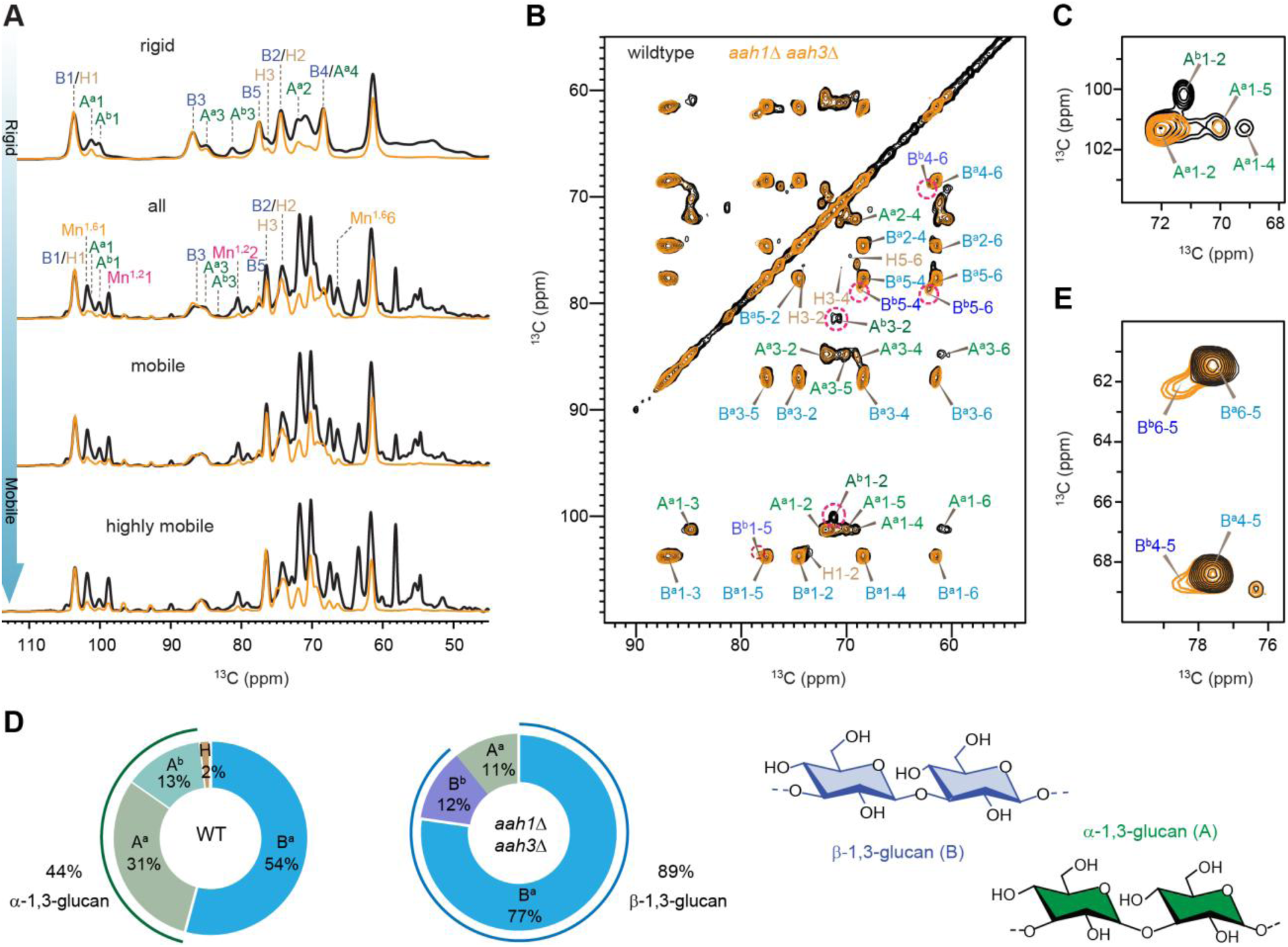
Solid-state NMR analysis of rigid polysaccharides in *S. pombe* cell walls. (**A**) 1D ^13^C spectra selectively detecting molecules with different dynamics in wildtype cells (black) and *aah1*Δ *aah3*Δ mutant (yellow). From top to bottom are ^13^C CP spectrum selecting rigid polysaccharides, ^13^C DP spectrum with long recycle delays quantitatively detecting all molecules, ^13^C DP spectrum with short recycle delays selecting mobile molecules, and ^13^C refocused INEPT selecting highly mobile molecules. Abbreviations: B, β-1,3-glucan; H, β-1,6-glucan; A, α-1,3- glucan; Mn^1,2^, α-1-2-mannose in mannan; Mn^1,6^, α-1,6-mannose in mannan. Superscript indicate structural subtypes within each polysaccharide, e.g., A^a^ and A^b^ indicate type-a and type-b α-1,3- glucans, respectively. (**B**) 2D ^13^C-^13^C 53-ms CORD spectrum resolving all carbons sites of rigid polysaccharides in the cell walls of wildtype (black) and *aah1*Δ *aah3*Δ (yellow). Predominant signals were observed for β-1,3-glucan (B) and α-1,3-glucan (A). Dashed line circles highlight the major differences between the two samples, with zoomed-in views provided in the later panels of this figure. (**C**) Wildtype sample contains strong signals of two forms of α-1,3-glucans (A^a^ and A^b^). A^a^ signals become weaker in the mutant, and A^b^ disappeared in the mutant. (**D**) Molar composition of rigid cell wall polysaccharides estimated using peak volumes in 2D ^13^C-^13^C 53-ms CORD spectra. Simplified structures of glucans are shown on the side. (**E**) Presence of two polymorphic forms of β-1,3 glucan (B^a^ and B^b^) in the *aah1*Δ *aah3*Δ mutant, while B^b^ is absent in the wildtype strain.

The rigid domain, critical for the mechanical integrity of the cell wall, was examined in detail using 2D ^13^C-^13^C correlation experiments. The analyses resolved all carbon sites in two allomorphic forms of β-1,3-glucans and two structural variants of α-1,3-glucans, which constitute the exclusive components of the rigid phase (**Fig. 4B**). We also confirmed the complete absence of chitin in this fungus (**Fig. S3**), consistent with other studies (15, 16). In the wildtype sample, strong signals corresponding to type-a polymorph and type-b polymorph of α-1,3-glucans were detected, as evidenced by their C1-C2 cross peaks (A^a^1-2 and A^b^1-2, **Fig. 4C**). Notably, type-b α- 1,3-glucans were entirely absent in the *aah1*Δ *aah3*Δ mutant, consistent with 1D spectral observations (**Fig. 4A-C**). Intensity analysis also revealed a significant decrease in type-a α-1,3- glucan content, from 31% in the wildtype to 11% in the mutant (**Fig. 4D**). Consequently, the overall α-1,3-glucan content declined markedly, from 44% in wildtype rigid cell walls to 11% in the *aah1*Δ *aah3*Δ mutant.

The substantial reduction in α-1,3-glucan content in *aah1*Δ *aah3*Δ cells and their phenotype were reminiscent of cells with defective *ags1* function, specifically the commonly used *ags1-664* mutant (46). We therefore wondered whether the mutation within *ags1-664*, which had not previously precisely determined, resided within the Ags1 GH13 domain (32, 33). Indeed, sequencing of *ags1- 664* revealed Y256F and R408H substitutions within the GH13 domain (**Fig. S4A**), underscoring its essential function. Given that *ags1, aah1*, and *aah3* all encode GH13 domain-containing proteins, we tested whether there were negative genetic interactions between *ags1-664* and the ɑ- amylase defective cells. As predicted for genes in a common pathway, a strong negative genetic interaction was observed between *aah3*Δ and *ags1-664* (**Fig. S4B**). To test if the Ags1 GH13 domain and Aah3 GH13 domains are functionally redundant, we over-produced Aah3 or Ags1 in *aah1*Δ *aah3*Δ cells. While overproduced Aah3 rescued the morphological defects of *aah1*Δ *aah3*Δ cells, over-production of Ags1 did not and overproduced Aah3 did not rescue *ags1-664* cells (**Fig. S4C, D**). We conclude that Aah1 and Aah3 enzymes are not functionally redundant with Ags1, and cells require both types of GH13 domain enzymes to support α-glucan chain assembly.

The weak β-1,6-glucan signals observed in the rigid fraction of the wildtype cell wall such as H3- 4 and H5-6 cross peaks (**Fig. 4B**), which accounted for only 2% (**Fig. 4D**), were absent in the *aah1*Δ *aah3*Δ mutant. This molecule was primarily detected in the mobile phase, as indicated by the strong carbon 3 (H3) peak at 76.5 ppm in the 1D ^13^C spectra that captured either mobile molecules or all molecules (**Fig. 4A**) and further supported by 2D spectra in subsequent sections. These findings suggest that a minor fraction of β-1,6-glucan, initially colocalized with rigid α-1,3- glucans and partially rigidified in the wildtype sample, shifted entirely to the mobile phase upon the depletion of α-1,3-glucans in the *aah1*Δ *aah3*Δ mutant.

This substantial loss of α-1,3-glucans in *aah1*Δ *aah3*Δ cell walls was accompanied by the emergence of a novel polymorphic form of β-1,3-glucans. The mutant retained the signals of type- a β-1,3-glucans (e.g., B^a^4-5 and B^a^6-5 cross peaks) and exhibited additional signals corresponding to a new structural variant (e.g., B^b^4-5 and B^b^6-5 cross peaks, **Fig. 4E**). The total β-1,3-glucan content increased from 55% in the wildtype to 89% in the mutant (**Fig. 4D**), consistent with the more intense Calcofluor White septum staining observed in *aah1*Δ *aah3*Δ cells compared to wildtype cells (**Fig. S4E)**. If the elevated level of β-1,3-glucan content in *aah1*Δ *aah3*Δ cells was a compensatory adaptation to the dramatic reduction in α-glucan content, *aah3*Δ cells would be expected to rely on β-1,3-glucan synthetase function. Indeed, *aah3*Δ cells had a mild negative genetic interaction with an allele of *bgs4 (cwg1-1)* and a strong negative genetic interaction with an allele of *bgs1 (cps1-191)* (**Fig. S4B**).

The Cell Integrity Pathway (CIP) (13, 60), a signaling cascade involving the protein kinase C homologs, Pck1 and Pck2, and the mitogen-activated protein kinase homolog, Pmk1, increases glucan synthase activity and gene expression when *S. pombe* cells sense defects in their cell walls (32, 61, 62). To test if the upregulation of β-glucans in *aah1*Δ *aah3*Δ cells could be through CIP signaling, we combined *pck2*Δ or *pmk1*Δ with *aah1*Δ or *aah3Δ,* crossed these strains with each other, and analyzed the resultant colonies after tetrad dissection. We found that *pck2Δ aah1*Δ *aah3*Δ and *pmk1Δ aah1*Δ *aah3*Δ combinations were synthetically lethal (**Fig. S4F**). These data suggest that there is a requirement for sufficient β-glucan synthesis mediated by the CIP when the α-glucan network is disrupted.

### *aah1*Δ *aah3*Δ cells display an extensively modified mobile domain

The mobile domain was confirmed to contain β-glucans, mannan, and α-glucans in both wildtype and mutant cells (**Fig. 5A**). Notably, the α-1,3-glucan in the mobile domain displayed greater polymorphism compared to the rigid core, with three polymorphic forms observed in wildtype cells, which were all largely absent in the *aah1*Δ *aah3*Δ mutant (**Fig. 5B**). Weak signals corresponding to α-1,4-linked glucopyranose residues (A^1,4^), representing minor structural domains within α-glucans, were detected in wildtype cells but were completely absent in the *aah1*Δ *aah3*Δ mutant, indicating a loss of these structural components (**Fig. 5C**). Intensity analyses demonstrated that the proportion of α-glucans in the mobile phase decreased from 33% in wildtype cell walls to 10% in the *aah1*Δ *aah3*Δ mutant (**Fig. 5D**). This indicates that the deletion of *aah1* and *aah3* genes led to a significantly simplified structure of the α-glucan matrix by eliminating the α-1,4-linked structural domain, concurrently reducing the polymorphism of the predominant α- 1,3-linked domains, and ultimately decreasing the total α-glucan content by threefold.

**Figure 5.**
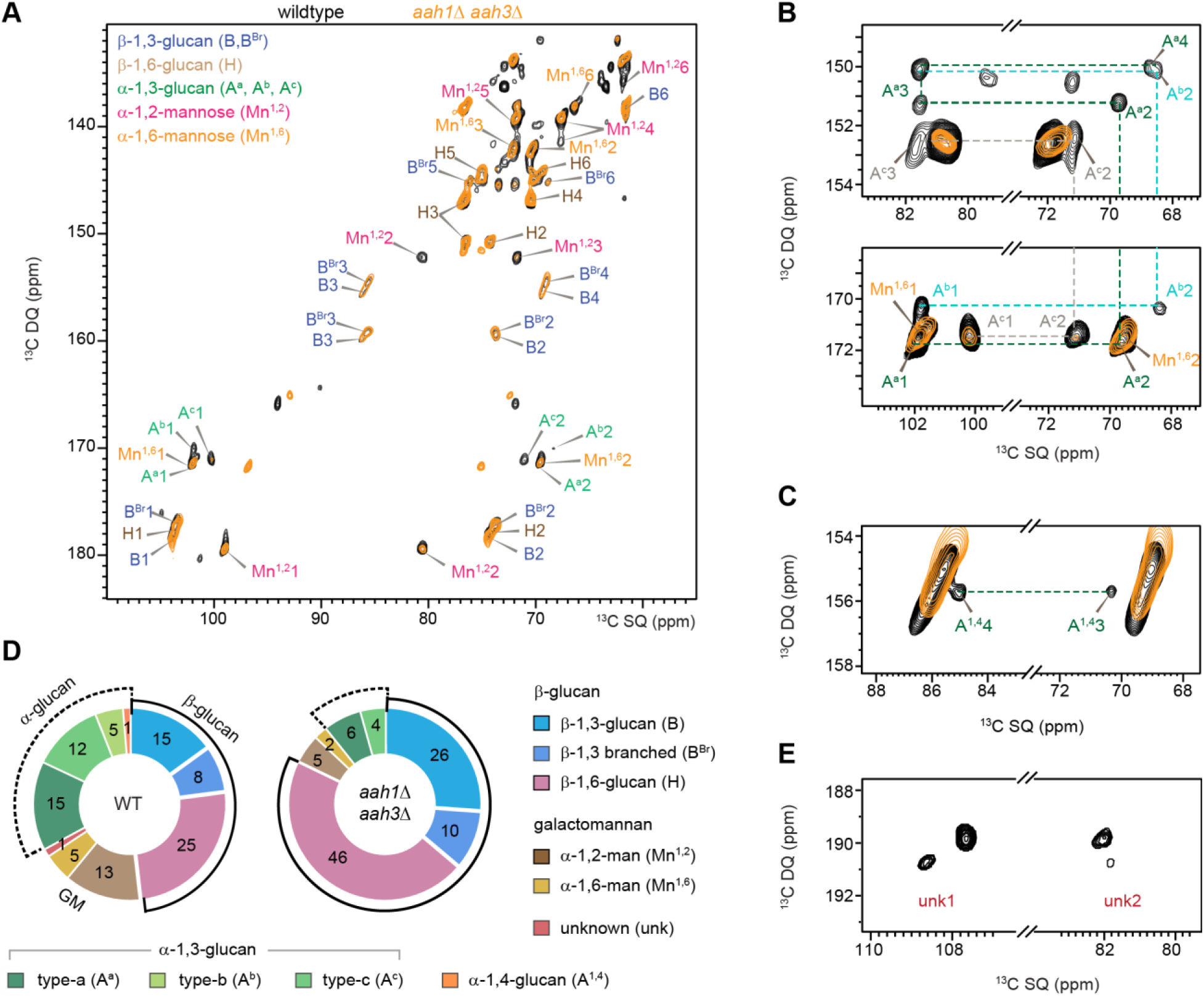
Mobile carbohydrate components in *S. pombe.* (**A**) 2D ^13^C DP refocused J-INADEQUATE spectrum showing signals of α-1,3-glucan (A), linear β-1,6-glucan (H), linear β- 1,3-glucan (B), β-1,3,6-glucopyranose unit (B^Br^) of the branched glucan, as well as α-1,6-mannan (Mn^1,6^) and α-1,2-mannose (Mn^1,2^) residues of mannan. (**B**) Three forms of α-1,3-glucan (A^a^, A^b^, A^c^) resolved in the mobile region of wildtype *S. pombe* (black), but significantly lower in the mutant. (**C**) Signals of α-1,4-glucan (A^1,4^) observed in wildtype sample but depleted in the double mutant. (**D**) Molar composition of mobile carbohydrates estimated using peak volumes in refocused J-INADEQUATE spectra. All the values presented in the plots are molar percentages (%). (**E**) Signal of unknown 1 (unk1) and unknown 2 (unk2) residues in the wildtype strain whereas it is absent in the *aah1*Δ *aah3*Δ mutant (yellow).

The β-glucan matrix displayed signals corresponding to linear β-1,3-glucan, linear β-1,6-glucan, and their branching sites containing β-1,3,6-glucopyranose units (**Fig. 5A**), which collectively accounted for 48% of mobile molecules in wildtype cell walls, increasing to 82% in the *aah1*Δ *aah3*Δ mutant (**Fig. 5D**). The quantities of both β-1,3-glucan and β-1,6-glucan nearly doubled in the mutant compared to wildtype cells. This observation aligns with findings in the rigid phase, as shown earlier in **Figure 4D**, although the dominant component shifted from β-1,3-glucan in the rigid phase to β-1,6-glucan in the mobile phase. These results collectively confirm a compensatory adaptation in the cell wall, independent of molecular dynamics, marked by upregulated β-glucan synthesis in response to the depletion of most α-glucans caused by gene deletion.

The galactomannan in the cell wall of both samples consistently contained α-1,6- and α-1,2-linked mannose residues as the major components; however, its abundance within the mobile cell wall fractions decreased from 18% in the wildtype sample to 7% in the mutant (**Fig. 5D**). Concurrently, the protein content within the mobile fraction exhibited a significant reduction in the mutant, a change that may include alterations in proteins associated with galactomannans (**Figs. S8, S9**). Earlier studies reported galactomannan as α-1,6-mannan with branches formed by α-1,2-mannose and terminal D-galactopyranose (Gal*p*) residues at non-reducing ends (19–24). However, in the present study, no detectable galactopyranose signals were observed, despite their high abundance in the solid-state NMR spectra of other fungal species.

We identified distinct NMR signals at 108 ppm and 82 ppm, corresponding to carbons 1 and 2, respectively, which match the characteristic chemical shifts of galactofuranose (Gal*f*) in the sidechains of galactomannan specific to *Aspergillus* species, while these signals were absent in the galactomannan-deficient mutant of *A. fumigatus* (63, 64). At first we suspected that the two forms of galactose units might coexist in equilibrium, allowing Gal*p* to convert to Gal*f*; however, *S. pombe* lacks galactopyranose mutase, the enzyme required for this conversion, ruling out this possibility (65). Upon further analysis, we considered that other five-membered monosaccharide units might also contribute to the observed NMR signals. One potential candidate is fructofuranose, whose biosynthesis is regulated by the proteins encoded by the *inv1* and *inv2* genes in *S. pombe* (66, 67). Its precise structural characteristics and role in the extracellular matrix remain to be fully elucidated. As this component constitutes less than 1% of the mobile carbohydrates in wildtype *S. pombe* and is undetectable in the mutant, with minimal effect on the compositional analysis, we set it aside as an unidentified sugar.

### *aah1*Δ *aah3*Δ mutants have a more rigid and less dynamic cell wall compared to wildtype

The molecular motions of the cell wall polysaccharides were investigated using ^1^H-T_1ρ_ relaxation, which was significantly slower in the *aah1*Δ *aah3*Δ mutant compared to wildtype cells for both β- 1,3-glucan and α-1,3-glucan (**Fig. 6A**). Fitting the data to a double-exponential equation revealed that the time constants of the long components increased from 4-5 ms in wild-type cells to 17-23 ms in the mutant for both β-1,3-glucan and α-1,3-glucan (**Fig. 6B**). Simultaneously, their relative fractions increased from 33-53% in wild-type cells to 75-86% in the mutant (**Fig. 6C**). The predominance of the slow-relaxation component, characterized by its increased proportion and prolonged relaxation time, indicates a substantial reduction in molecular dynamics on the microsecond timescale in the mutant. Fitting the data to a single-exponential equation gave the same trend: the average ^1^H-T_1ρ_ relaxation time constant in the mutant were an order of magnitude longer than in the wildtype cells, increasing from 2.2 ms to 13.6 ms for β-1,3-glucan and from 2.2 ms to 11.5 ms for α-1,3-glucan (**Fig. 6D** and **Fig. S5**). Consistently, moderate increases were observed in ^13^C-T_1_ relaxation times, rising from 1.2 s to 1.4 s for β-1,3-glucan and from 2.1 s to 2.9 s for α-1,3-glucan, suggesting reduced local dynamics on the nanosecond timescale (**Fig. S6**).

**Figure 6.**
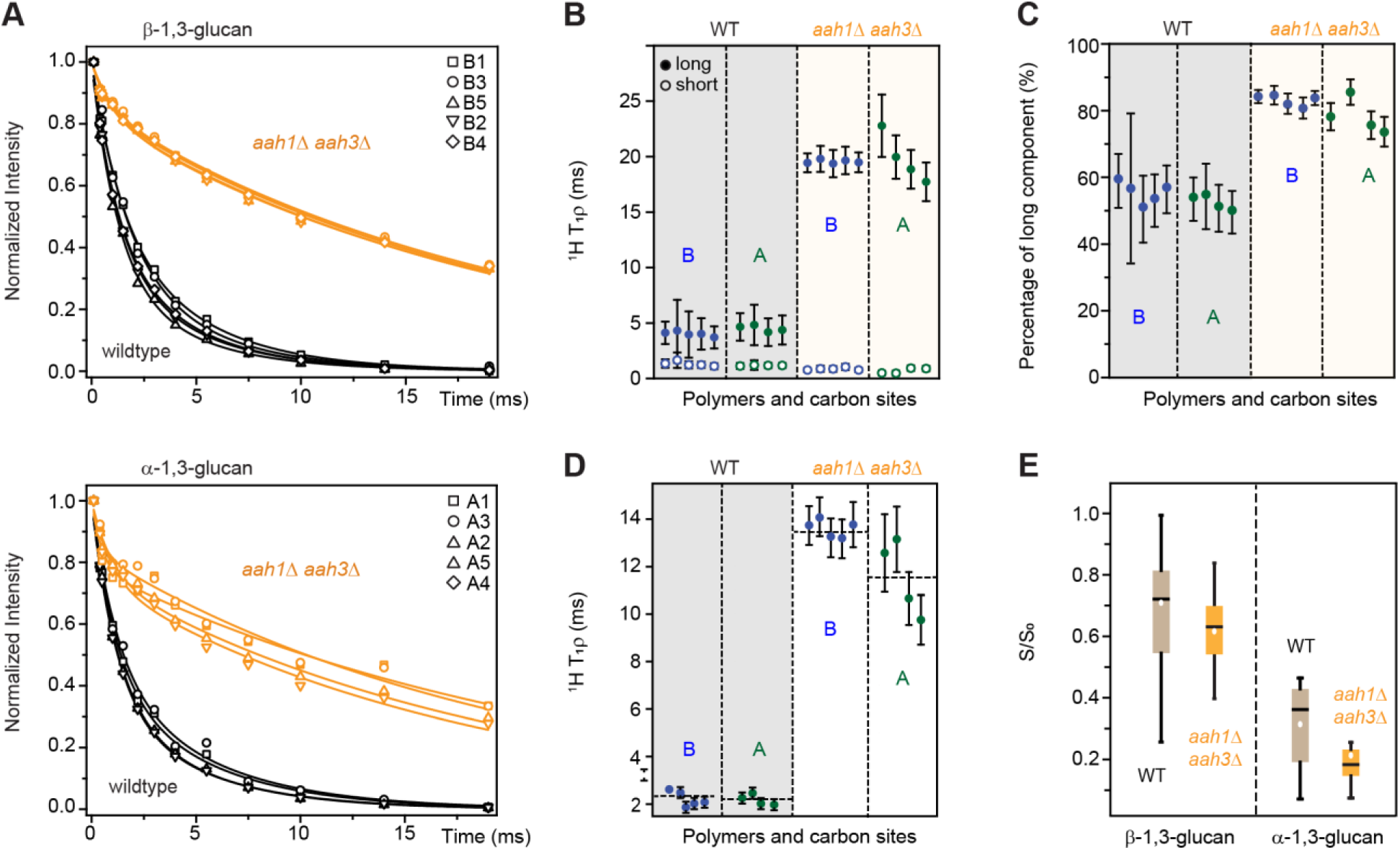
FNMR relaxation and water-edited intensities of polysaccharides. (**A**) ^1^H-T_1ρ_ relaxation curves fit to double-exponential equations were plotted for different carbon sites in β- 1,3-glucan (top) and α-1,3-glucan (bottom) as comparisons between wildtype cells (black) and *aah1*Δ *aah3*Δ mutant (yellow). (**B**) ^1^H-T_1ρ_ relaxation time constants from double-exponential fitting for different carbon sites of β-1,3-glucan (blue) and α-1,3-glucan (green) in the wildtype (left; shaded in grey) and *aah1*Δ *aah3*Δ mutant (right; shaded in pale yellow). Long and short components of ^1^H-T_1ρ_ relaxation are represented using filled and open circles, respectively. (**C**) Fraction (%) of the long-component of ^1^H-T_1ρ_ relaxation. (**D**) ^1^H-T_1ρ_ relaxation time constants from single-exponential fitting. (**E**) Box-and-whisker diagram plotting the relative water-edited intensities (S/S_0_) of β-1,3-glucan and α-1,3-glucan in wildtype cells (grey) and *aah1*Δ *aah3*Δ mutant (yellow). For each sample, n=36 for β-1,3-glucan and n=12 for α-1,3-glucan. The boxes represent the IQR, with whiskers extending to 1.5 IQR. Mean values are represented by open circles and medians by horizontal lines

Meanwhile, water retention in these glucans, assessed by water-edited S/S_0_ intensities at different carbon sites (68–70), was also lower in the mutant than in wildtype cells, with average values dropping from 0.71 to 0.61 for β-1,3-glucan and from 0.51 to 0.20 for α-1,3-glucan (**Fig. 6E** and **Fig. S7**); therefore, polymer dynamics on the microsecond timescale were impeded as well. Therefore, the thick and abnormally shaped cell walls of the *aah1*Δ *aah3*Δ mutant have reduced water activity and accessibility within the wall, and substantially restricted the collective motions of carbohydrates on microsecond timescale and the rapid local motions on the nanosecond timescale, resulting in markedly rigidified cell walls.

## DISCUSSION

In this study, we applied solid-state NMR spectroscopy to analyze the cell wall composition and structure of intact fission yeast cells. We determined that the loss of two GPI-anchored ɑ-amylase homologs, Aah1 and Aah3, resulted in a dramatic reduction of cell wall α-1,3-glucans and galactomannans with a compensatory increase in β-1,3-glucans. Indeed, TEM analysis showed that many *aah1*Δ *aah3*Δ cells lack the outermost galactomannan cell wall layer, and the overall cell wall becomes thicker than in wildtype cells. As a result of these cell wall abnormalities, cell shape and cell separation following cytokinesis of *aah1*Δ *aah3*Δ cells are significantly disrupted leading to the formation of clumps of spheroid cells. Thus, not only does rod-shaped cell morphology of *S. pombe* depend on α-glucan synthase function but it also depends on polysaccharide modifying enzymes positioned outside the cellular environment that coordinate with α-glucan synthase activities.

Solid-state NMR allows for the direct characterization of polysaccharide composition and the physical properties of intact cells, leading to conceptual advances in understanding cell wall architecture across a diverse range of fungal species, including *Aspergillus* (64, 71–74), *Cryptococcus* (75–78), *Candida* (79), *Schizophyllum* (80–82), as well as *Rhizopus* and *Mucor* species (83). The application of this technique to *S. pombe* in this study uncovered several novel findings, including insights into the distinct structural roles of various polysaccharides in cell wall construction. It is now understood that the rigid core of *S. pombe* mainly comprises two forms of linear α-1,3-glucan and a type of β-1,3-glucan, the latter playing a crucial role in regulating water activity (**Fig. 7A**). These chemically identical polymorphic glucan forms may be distinct due to differences such as conformation, hydrogen bonding or binding partners. These molecules also extend into the dynamic matrix supporting the mechanical scaffold, alongside contributions from galactomannan, most β-1,6-glucans, a third type of linear α-1,3-glucan, and α-1,4-linked domains of α-glucans. Overall, our findings are consistent with previous studies that determined the composition of purified *S. pombe* cell wall material using various methods (84–88), while the spatial information obtained aligns with results from techniques such as immuno-electron microscopy (16, 17, 84, 89), and simultaneously provides novel insights into the structural polymorphism of carbohydrates in intact *S. pombe* cells.

**Figure 7.**
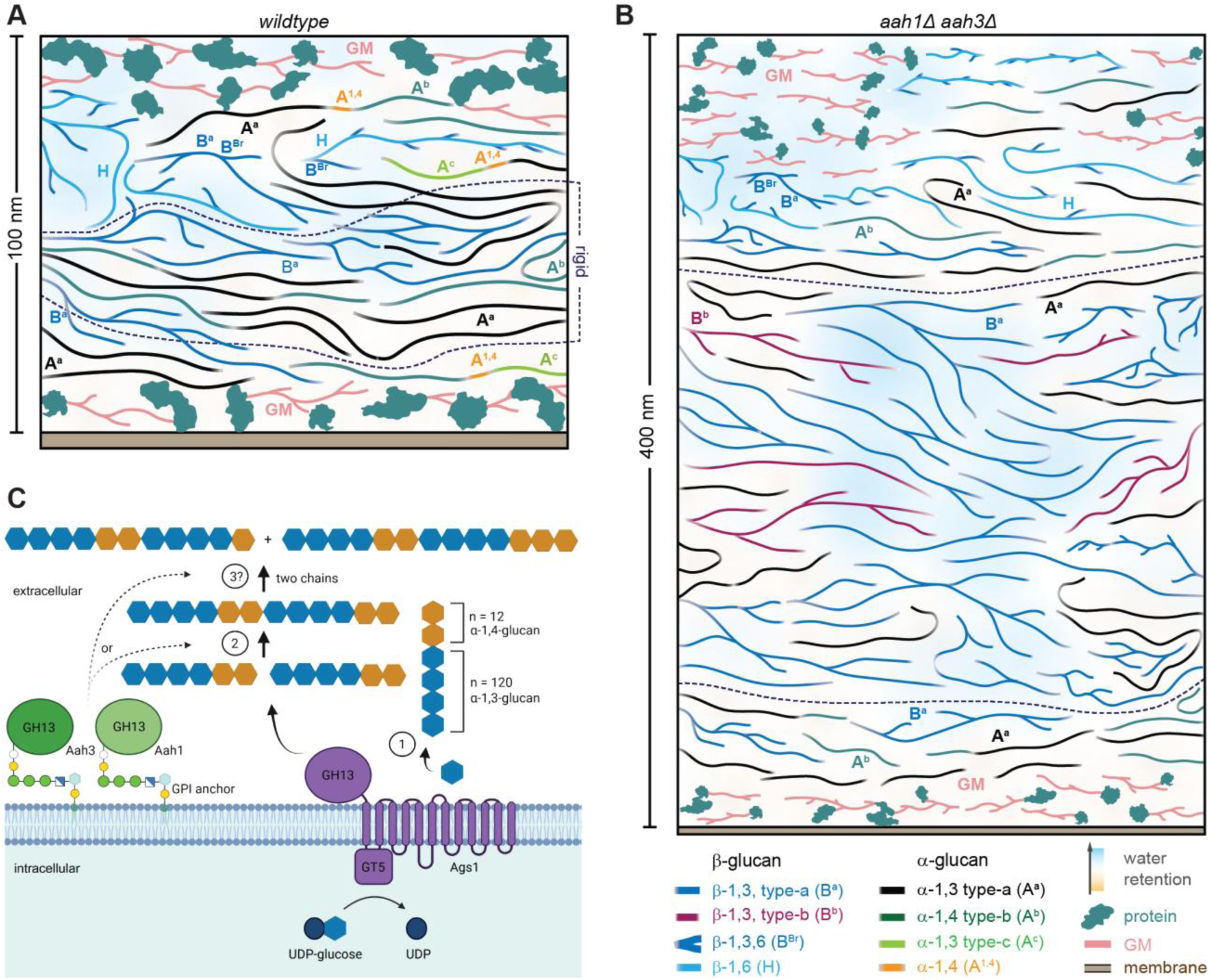
Proposed function of Aah3 and Aah1 in collaboration with Ags1. Illustrative cartoons depict simplified *S. pombe* cell wall structures for (**A**) wildtype and (**B**) *aah1*Δ *ahh3*Δ mutant cells, with the latter shown at 50% of the length scale used for the former. The legends illustrate four types of β-glucans, four types of α-glucans, galactomannan (GM), membranes, and proteins. The background color uses light blue to represent the possible hydration profile. In each panel, the region between the two dashed lines represents the rigid core, while the surrounding areas are mobile. The molecular composition within the rigid and mobile fractions is represented in the plots, though not necessarily to scale. The illustrations integrate cell wall thickness from imaging, along with composition, structural polymorphism, and dynamic and hydration profiles informed by ssNMR, as well as previous biochemical data and models of *S. pombe* cell walls. (**C**) Schematic representation of biosynthesis and transglycolysis of α-1,3-glucan in *S. pombe*. The existing model of Ags1 function in *S. pombe* (14) hypothesizes that the Ags1 (purple) synthase/GT5 domain catalyzes the addition of α-1,3-glucan (blue) to an α-1,4-glucan primer (orange) (step 1). Subsequently, two chains are joined by the extracellular GH13 domain of Ags1 (step 2). Aah1 and Aah3 (light and dark green, respectively) may have the same enzymatic specificity as the Ags1 GH13 domain, but their unique spatial determinants could underlie their distinct functional requirement (step 2). Another possibility is that Aah1 and Aah3 perform a disproportionation reaction downstream of Ags1 (step 3).

Comparative NMR analysis of the wildtype and *aah1*Δ *aah3*Δ mutant cells revealed a significant reduction in α-glucan content in *aah1*Δ *aah3*Δ cells, complete depletion of one α-1,3-glucan form from both rigid and mobile fractions, partial loss of the other α-1,3-glucan form within the rigid domain, and the complete absence of all α-1,4-glucans from the mobile domain (**Fig. 7B**). In contrast, β-glucan content increased, accompanied by the emergence of a new form of rigid β- glucan not present in wildtype cells (**Fig. 4E**), along with a two-fold increase in the levels of β- 1,6- and β-1,3-glucans in the mobile phase (**Fig. 5D**). The spatial organization of these molecules is reshuffled in the *aah1*Δ *aah3*Δ mutant, resulting in thickened, rigidified, and more hydrophobic cell walls (**Fig. 3B** and **Fig. 6**). Our genetic data suggest that the increased β-glucan content of *aah1*Δ *aah3*Δ cell walls is required for their viability and that the increase is achieved via CIP activity. These observations provide evidence for a feedback mechanism in which upregulation of β-glucan synthases partially compensates for disturbances in the ɑ-glucan network. A reciprocal result indicating this feedback mechanism exists is that over-production of Ags1 results in reduced β-glucan content (32). Elucidating the mechanism of these compensatory relationships will be a fruitful area of future study.

Another outstanding question from the current study is how changes in cell wall glucan content result in changes to galactomannan content and organization. The same loss of galactomannans we observed for *aah1Δ ahh3*Δ cells was also observed in *ags1* and *bgs4* deficient cells, in addition to cells treated with a β-glucan synthase inhibitor, papulacandin B (24, 46, 90, 91). Conversely, over-production of *ags1* results in a 2-fold increase in cell wall galactomannan content (32). It was previously suggested that mannose residues may associate with α-glucan fibrils, although it remains unclear how these networks are coordinated (46). Certainly, if the galactomannans depend on a connection with α-glucan fibrils to form the outer cell wall layer, a drastic reduction in α- glucan fibrils could explain the cell wall shedding phenotype observed in *aah1Δ ahh3*Δ and *ags1* defective cells (46). The shedding might also reflect the inherently dynamic nature of fungal cell walls, where the depletion of a single structural polysaccharide can trigger a complete reshuffling of biosynthetic pathways for other carbohydrates, resulting in a fully restructured cell wall (64, 92).

Our analysis of Aah1 and Aah3 using genetics, imaging and solid-state NMR allowed us to incorporate these enzymes into the current model of α-glucan synthesis in *S. pombe* (**Fig. 7C**). Given that *aah1*Δ *aah3*Δ cells are nearly indistinguishable from cells with defective Ags1 GH13 function (46), we propose that Aah1 and Aah3 are necessary for the biosynthesis of α-glucan chains. Whether Aah1/Aah3 act on the same substrates as Ags1 or at a distinct step of ɑ-glucan chain elongation remains to be determined. However, our results indicating that Aah1/Aah3 and Ags1 function non-redundantly suggest that they have unique biochemical activities or that spatial differences may impart their distinct functional requirement (**Fig. 7C**). Although the exact catalytic activity of Aah1/Aah3 has yet to be established, transglycosylation is a common activity of GH13 domain containing enzymes (93–95) and by analogy and homology to *Aspergillus niger* GPI- anchored ɑ-amylase/GH13 enzymes, we propose Aah1 and Aah3 are transglycosylases. GH13 domains exhibiting hydrolase activity contain a histidine in their active site (96). In contrast, the *A. niger* GPI-anchored ɑ-amylase/GH13 enzymes lack the active site histidine required for hydrolase activity and exhibit a 4-α-glucanotransferase activity in which α-glucan units from one chain are removed and added to another, generating one shorter and one longer α-glucan chain (96). This reaction is referred to as a disproportionation reaction and has been observed for additional ɑ-amylase/GH13 enzymes (94). Because Aah1 and Aah3 also lack a histidine in the active site, it is plausible that they act downstream of Ags1 to modify ɑ-glucan chain architecture.

GH13 domains exist in twelve *S. pombe* proteins, of which all but one belongs to the α-glucan synthase or α-amylase family (35). While we have analyzed the four α-amylases that are GPI-anchored and expressed during vegetative growth, three additional α-amylases are present in *S. pombe* including one that is expressed during meiosis and two that are not GPI-anchored (35). Thus, α-amylases may contribute to cell wall construction during other cell-cycle stages, and they may also modify the cell wall when not anchored to the outer leaflet of the plasma membrane. Further, in addition to Aah1 and Aah3, *S. pombe* contains 55 additional GPI-anchored proteins of which 7 are implicated in carbohydrate catabolism (35). While Ags1 contains intrinsic cell wall modifying activity in addition to synthase activity, β-glucan synthases contain only synthase activity. Whether other cell wall modifying proteins link to β-glucan synthases, like the Sbg1 (97, 98), to coordinate the remodeling of the cell wall remains to be determined. Further studies on the functions of these proteins may provide insights into how cells construct a wall that achieves the precise balance between flexibility and rigidity.

## METHODS

### Preparation of ^13^C, ^15^N yeast samples

*S. pombe* strains (**Table S1**) were grown in yeast extract (YE) medium(99) or Edinburg minimal medial (EMM) with appropriate supplements (99). Strains were constructed using standard *S. pombe* mating, sporulation, and tetrad dissection techniques (100). For solid-state NMR analysis, *S. pombe* wildtype and mutant strains (*aah1*Δ *aah3*Δ) were inoculated into 250-mL Erlenmeyer flasks containing 100 mL of the autoclaved ^13^C/^15^N-labeled growth medium and incubated at 30°C with constant shaking at 200 rpm (Corning LSE Shaking Incubator, 6753). The cultures were grown to an optical density (OD) of 0.5-1.0. Cells were harvested by centrifugation at 5000 × g for 20 minutes (Thermo Fisher Scientific, 75016014). The resulting cell pellets were washed thoroughly with Milli-Q water to remove small water-soluble molecules and excess ions, ensuring retention of natural cellular hydration. Approximately 45–50 mg of the prepared cell material was packed into 3.2 mm magic-angle spinning (MAS) rotors (HZ16916, Cortecnet) for subsequent solid-state NMR analysis.

The EMM media used for labeling *S. pombe* contains ^13^C-labeled D-glucose (20.0 g/L, CLM- 1396-PK, Cambridge Isotope Laboratories) and ^15^N-labeled ammonium chloride (5.0 g/L, NLM- 713-PK, Cambridge Isotope Laboratories), 3.0 g/L potassium hydrogen phthalate, 2.2 g/L sodium hydrogen phosphate, 20.0 mL/L salt solution (52.5 g/L magnesium dichloride hexahydrate, 0.735 g/L calcium dichloride dihydrate, 50.0 g/L potassium chloride, and 2.0 g/L sodium sulfate), 1.0 mL/L vitamin solution (1.0 g/L pantothenic acid, 10.0 g/L nicotinic acid, 10.0 g/L inositol, 10.0 mg/L biotin), 0.1 mL/L mineral solution (5.0 g/L boric acid, 4.0 g/L manganese sulfate, 4.0 g/L zinc sulfate heptahydrate, 2.0 g/L ferrous chloride hexahydrate, 0.4 g/L molybdic acid, 1.0 g/L potassium iodide, 10.0 g/L citric acid, and 0.4 g/L copper sulfate pentahydrate), along with 225 mg/L adenine, 225 mg/L uracil, and 225 mg/L leucine. The solution was autoclaved prior to use.

### TEM analysis of cell wall thickness and morphology

*S. pombe* strains were grown to mid-log phase at OD_595_ ∼0.3 and then pelleted and resuspended in molten 2% low-melt agar, 1% sucrose, 50 mM HEPES pH 7.3. The agar was cut into 1 mm cubes and rapidly frozen by plunge freezing into liquid ethane. Samples were freeze-substituted in 1.5% uranyl acetate in methanol at -90°C for 48 hours followed by infiltration HM20 Lowicryl for 48 hours at -45°C. The resin was UV polymerized at -45°C for 24 hours, followed by an additional 24 hours of UV at 0°C. The samples were cut on a Leica Enuity at a nominal thickness of 70 nm and collected on either 300 mesh Ni grids or 400 mesh carbon coated Cu grids. Grids were stained with 1% phosphotungstic acid, 2% uranyl acetate, and lead citrate to contrast the cell walls. Samples were imaged by transmission electron microscopy (TEM) on a JEOL 2100+ TEM operating at 200 keV using an AMT nanosprint 15mkII CMOS camera. Cell wall thickness was measured using ImageJ (software version V1.8.0_172).

### Solid-state NMR experiments

NMR experiments were conducted on two Bruker Avance Neo spectrometers at MSU Max T. Roger facility, with ^1^H Larmor frequencies of 800 MHz (18.8 Tesla) and 400 MHz (9.4 Tesla). All experiments were collected using two 3.2-mm MAS HCN probes under 15 kHz MAS at 293 K. ^13^C chemical shifts were externally referenced to the adamantane CH_2_ peak at 38.48 ppm on the tetramethyl silane (TMS) scale. Typical radiofrequency field strengths, unless specifically mentioned, were 80-100 kHz for ^1^H decoupling, 62.5 kHz for ^1^H hard pulses, and 50-62.5 kHz for ^13^C. The experimental parameters are summarized in **Table S2**.

ssNMR experiments were carried out using 1D and 2D experimental methods. 1D ^13^C spectra were measured using different polarization methods to selectively detect the rigid and mobile components (**Fig. S9**). The rigid components were detected by 1D ^13^C cross-polarization (CP) using 1-ms contact time (101, 102). Mobile components were measured by 1D ^13^C direct polarization (DP) using short (2 s) recycle delays. Extending the recycle delays to 35 s allowed us to achieve quantitative detection of all molecules through the same ^13^C DP experiment. The most mobile and highly solvated molecules are identified using a J-coupling based refocused INEPT experiments(103). Resonance assignment of rigid molecules was achieved through the measurement of 2D ^13^C-^13^C 53-ms CORD homonuclear correlation spectra (104), which detect intramolecular cross-peaks. For mobile molecules, 2D DP refocused J-INADEQUATE spectra were measured to track through bond connectivity, with each of the four tau delays optimized to 2.3-ms (105). The resolved chemical shifts were compared with the values indexed in the Complex Carbohydrate Magnetic Resonance Database (CCMRD) to validate the chemical nature of the carbohydrates (106). In addition to carbohydrate signals, protein signals were also detected, primarily in the mobile phase (**Fig. S8**). The ^13^C chemical shifts of carbohydrate and proteins identified in *S. pombe* have been tabulated in **Table S3**.

Water accessibility of polysaccharides was accessed using a water-edited 2D ^13^C-^13^C correlation experiment (68, 69). The experiment initiated with ^1^H excitation followed by a ^1^H-T_2_ filter (1.0- ms × 2 for the wildtype sample and 1.4-ms × 2 for the mutant), which eliminated 95% of carbohydrate signals but retained 70-80% of water magnetization (**Fig. S7**). The magnetization of water was then transferred to polysaccharide using a 4-ms ^1^H mixing period and then transferred to ^13^C through a 1-ms ^1^H-^13^C CP for ^13^C detection. A 50-ms DARR mixing period was used for both the water-edited spectrum and a control 2D spectrum showing full intensity. The ratio between the intensities of the water-edited spectrum (S) and the control spectrum (S_0_) was quantified; these S/S_0_ ratios reflect the relative extent of hydration at different carbon sites. The intensities were pre-normalized by the number of scans of each spectrum.

Polysaccharide dynamics were examined using ^13^C-detected ^1^H-T_1ρ_ relaxation, which was also measured using a Lee-Goldburg (LG) spinlock sequence combined with LG-CP for suppressing ^1^H spin diffusion during both the spinlock and the CP contact. This allowed site-specific examination of the ^1^H-T_1ρ_ relaxation of protons sites through detection of their directly bonded carbons. The decay of peak intensity was fit to both a single exponential function (**Table S4**) and a double exponential function (**Table S5**). In addition, ^13^C spin-lattice (T_1_) relaxation was measured, where the z-filter duration ranged from 0.1-μs to 12-s (**Fig. S6** and **Table S5**) (107). Each well-resolved peak was considered and the decay of its intensity was quantified as the z-filter time increased, and the data were fit to a single exponential equation to calculate the ^13^C-T_1_ relaxation time constant. The absolute intensity of each peak was pre-normalized by the number of scans. All the spectra were analyzed using Topspin 4.3.0 and all the graphs were generated through OriginPro 9.0. Illustrative figures were prepared using Adobe Illustrator 2023 V27.2.0.

The molecular composition was estimated by analyzing the intensities of distinct carbohydrate signals observed in two-dimensional ^13^C spectra (83). Rigid molecules were analyzed using CORD spectra, while mobile molecules were evaluated using DP refocused J-INADEQUATE spectra. The estimation was based on peak volumes, with only well-resolved signals included in the analysis. The relative abundance of each polysaccharide was quantified by normalizing the summation of integrated peak volumes by their respective cross-peak counts. The standard error for each polysaccharide was calculated by dividing the standard deviation of the integrated peak volumes by the total cross-peak count. The overall standard error was determined as the square root of the sum of the squared standard errors for all polysaccharides. To estimate the percentage error for each polysaccharide, the standard error of the specific polysaccharide was divided by its average integrated peak volume. This value was then multiplied by the ratio of the polysaccharide’s standard error to the total integrated peak volume, adjusted by its relative abundance.

### Fluorescence and DIC microscopy

Images were acquired with 1) a personal DeltaVision microscope system (Leica Microsystems), which includes an Olympus IX71 microscope, 60×NA 1.42 Plan Apochromat and 100×NA 1.40 U-Plan S-Apochromat objectives, live-cell and standard filter wheel sets, a pco.edge 4.2 sCMOS camera, and softWoRx imaging software or 2) a Zeiss Axio Observer inverted epifluorescence microscope with Zeiss 63X oil (1.46 NA) and captured using Zeiss ZEN 3.0 (Blue edition) software and Axiocam 503 monochrome camera (Zeiss).

### Molecular Biology

*ags1*^+^ with 300 bp of the 5’ UTR and 300 bp of the 3’ UTR was amplified from genomic DNA and cloned into the PstI restriction site of pITR2 using Gibson assembly (108, 109). *aah3^+^* was amplified from genomic DNA and cloned into the NdeI/BamHI sites of pREP81 using Gibson assembly (110). Whole plasmids were sequenced to confirm successful cloning by Plasmidsaurus using Oxford Nanopore Technology with custom analysis and annotation. Expression of *aah3* from the *nmt81* promoter was induced by switching cells from EMM containing 5 µM thiamine to EMM lacking thiamine for 24-38 hours prior to. Linear/amplicon sequencing of the first 3 kb of *ags1-664* was performed by Plasmidsaurus using Oxford Nanopore Technology with custom analysis and annotation.

## DATA AVAILABILITY

All relevant data that support the findings of this study are provided in the article and Supplementary Information. All the original ssNMR data files will be deposited in the Zenodo repository and the access code and DOI will be provided.

## AUTHOR CONTRIBUTIONS

A.J., M.E.H.E.N., and A.K.A.A. conducted solid-state NMR experiments and data analysis. A.H.W, M.G.I and L.A.T conducted imaging experiments and image analysis. A.H.W and M.G.I performed all yeast growth assays. A.J. and A.H.W. prepared fungal samples. All authors wrote the manuscript. K.L.G. and T.W. designed and supervised the project.

## ACKNOWLEDGMENT

The solid-state NMR research reported in this publication was supported by National Institutes of Health (NIH) under award number R01AI173270 to T.W. Sample preparation and imaging analysis were supported by NIH grant R35GM131799 to K.L.G. Transmission electron microscopy experiments were performed in part through the use of the Vanderbilt Cell Imaging Shared Resource (CISR) core (supported by NIH grants 1R24OD037694 and 1S10OD034315).

