## Supplementary Material for "Solid-State NMR Analysis of *Schizosaccharomyces pombe* Reveals Role of *α*-Amylase Family Enzymes in Cell Wall Structure and Function"

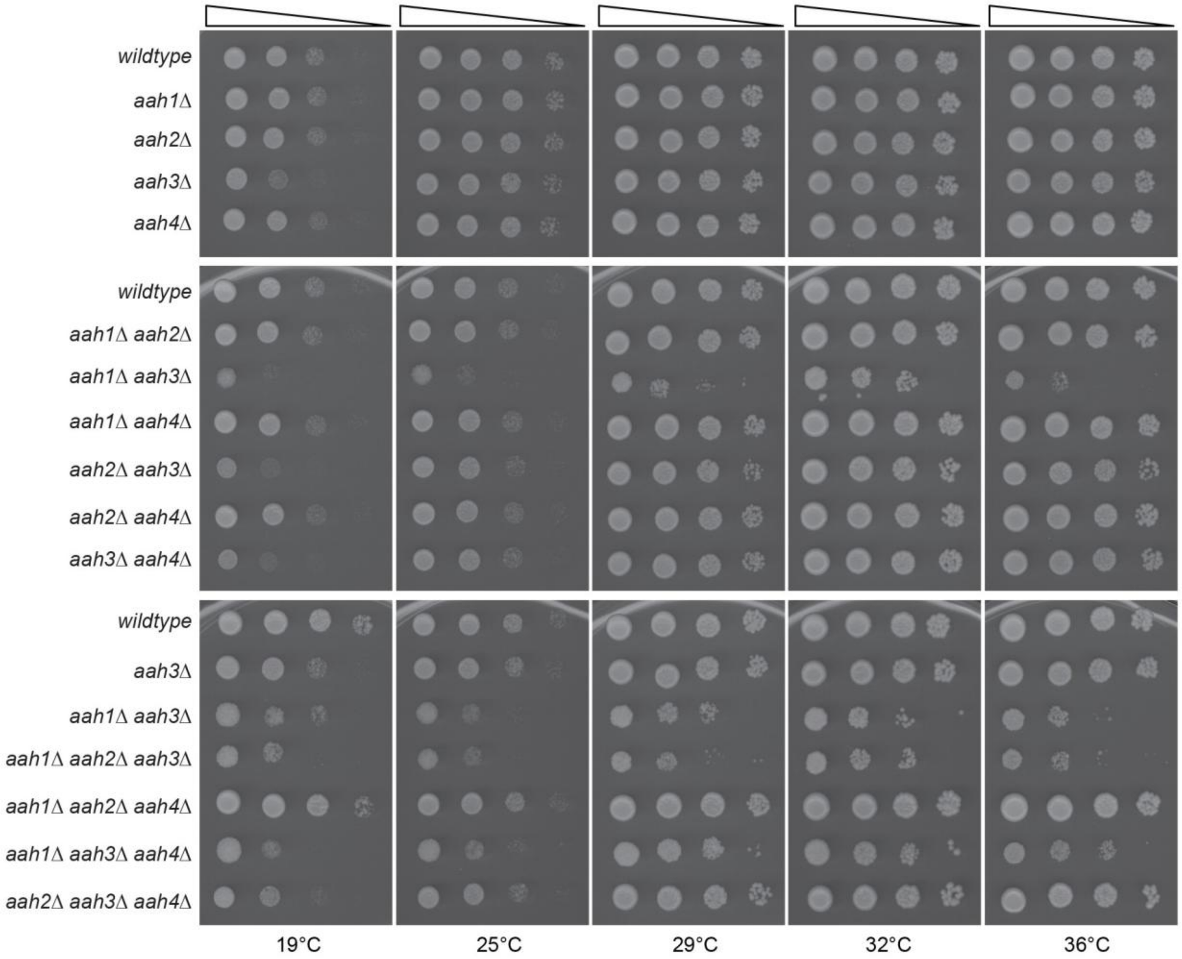


**Figure S1. Genetic analysis of cells lacking α-amylase encoding genes.** Serial 10-fold dilutions of the indicated strains spotted on YE plates and incubated at the indicated temperatures for 3-4 days prior to imaging.

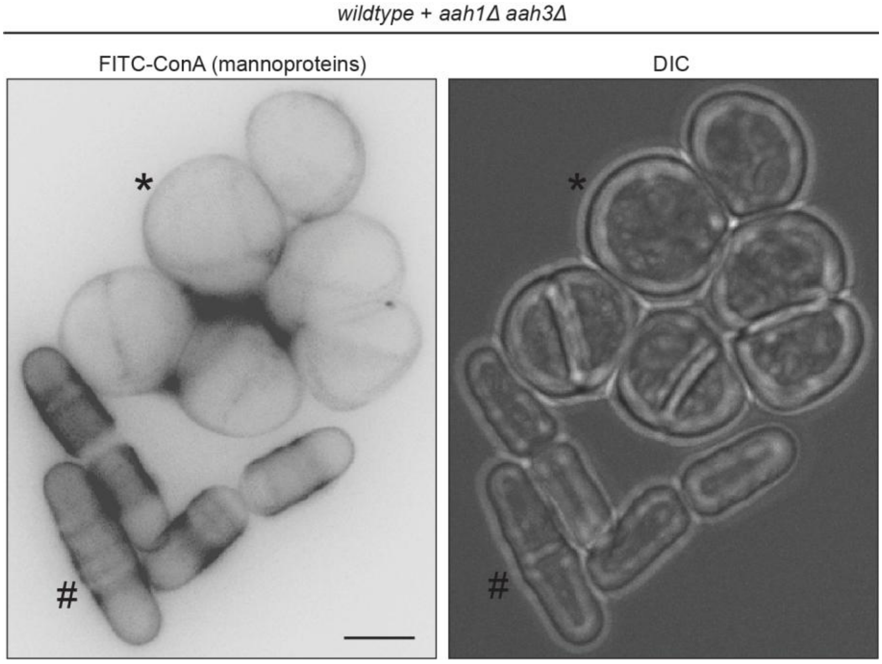


**Figure S2. *aah1Δ aah3Δ* cells have reduced galactomannan compared to wildtype cells.** Live-cell imaging of a 1:1 mixture of wildtype and *aah1Δ aah3Δ* cells stained with FITC-Concanavalin A (ConA). To distinguish each genotype, wildtype cells also expressed Sad1-mCherry (not shown). "#" mark wildtype cells and "*" mark a cluster of *aah1Δ aah3Δ* cells. Scale bar, 5 μm.

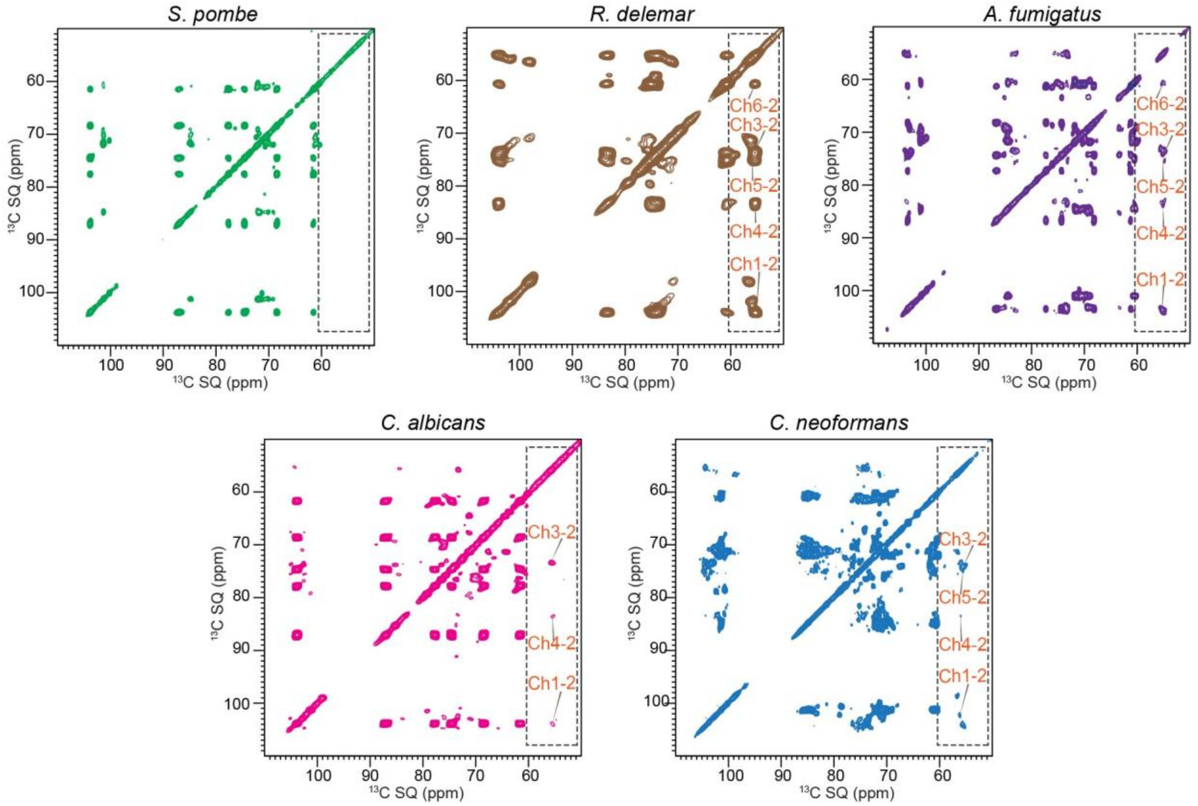


**Figure S3. Chitin signals from different fungal organisms.** 2D $^{13}$C-$^{13}$C 53-ms CORD spectra of different fungal organisms showing chitin as a major cell wall polysaccharide in their cell wall, interestingly the unicellular *S. pombe* does not show chitin signals from their cell wall. The *S. pombe* data was compared with those from *Rhizopus delemar* (a chitin-rich fungus), *Aspergillus fumigatus*, *Candida albicans*, and *Cryptococcus neoformans*. All the $^{13}$C-$^{13}$C 53-ms CORD spectra were measured on 800 MHz spectrometer at 15 kHz MAS**.**

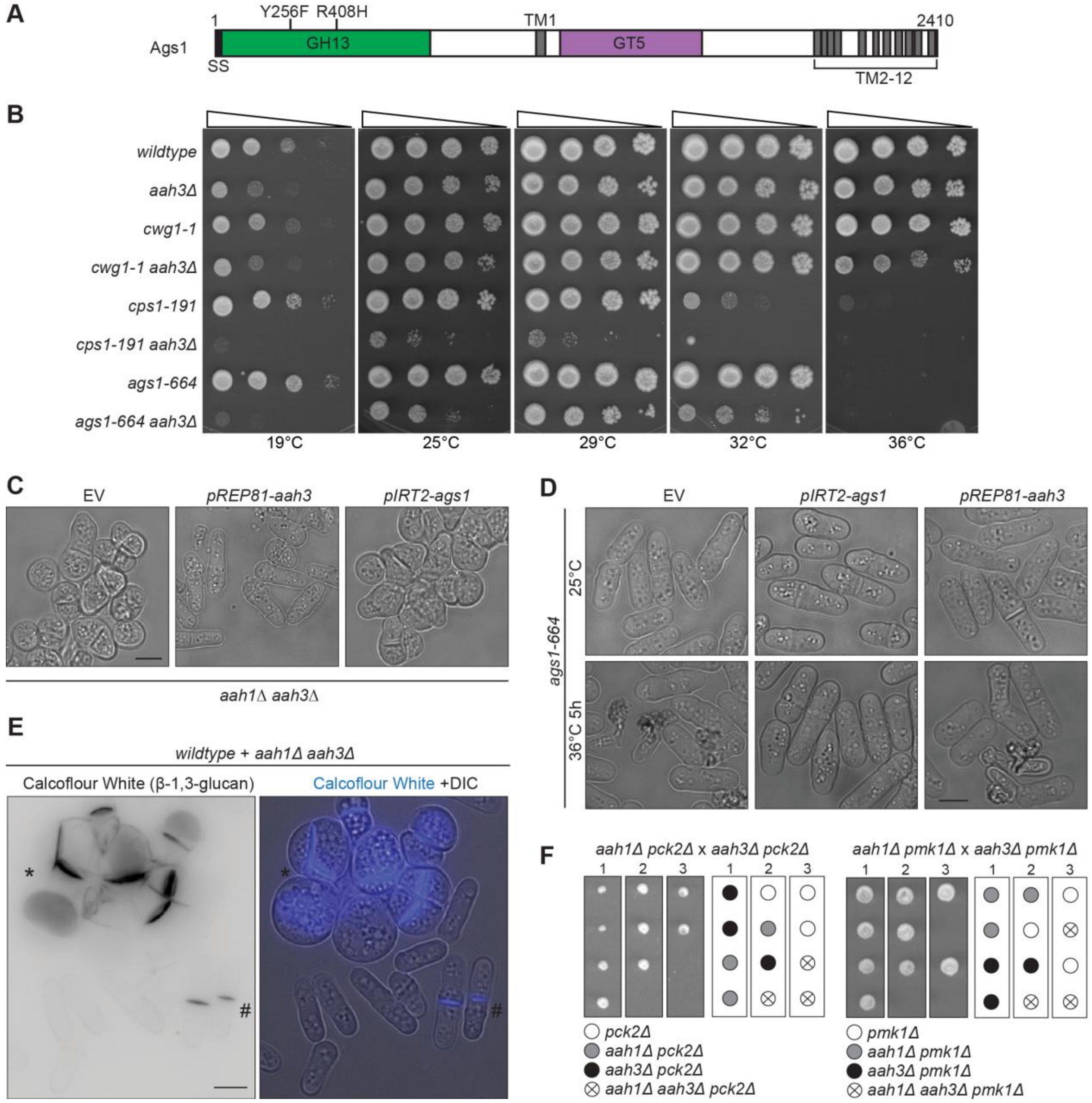


**Figure S4. Genetic analysis of *aah3*Δ cells**. (**A**) A schematic of Ags1, drawn to scale, with the domains and the *ags1-664* encoded mutations indicated. The glycoside hydrolase 13 (GH13), glycosyltransferase 5 (GT5) and transmembrane (TM) domains are shown. (**B**) Serial 10-fold dilutions of the indicated strains spotted on YE plates and incubated at the indicated temperatures for 3-4 days prior to imaging. (**C**) Live-cell DIC imaging of *aah1Δ aah3Δ* cells induced to express an empty vector (EV) or *aah3*$^{+}$ from the *nmt81* promoter for 24 hours prior to imaging. *ags1*$^{+}$ was constitutively expressed from its endogenous promoter within a pIRT2 vector. Cells were grown at 32°C. (**D**) Live-cell DIC imaging of *ags1-664* cells induced to express an empty vector (EV) or *aah3*$^{+}$ from the *nmt81* promoter for 24 hours prior to imaging. *ags1*$^{+}$ was constitutively expressed from its endogenous promoter within a pIRT2 vector. Cells were grown at 25°C and shifted to 36°C for 5 hours and imaged at both time points. (**E**) Live-cell imaging of a 1:1 mixture of wildtype and *aah1Δ aah3Δ* cells stained with 5 µg/ml of Calcofluor White. "#" mark wildtype cells and "*" mark a cluster of *aah1Δ aah3Δ* cells. Scale bars, 5 µm. **(F)** Representative tetrads and schematics of the analyses of the indicated genetic crosses.

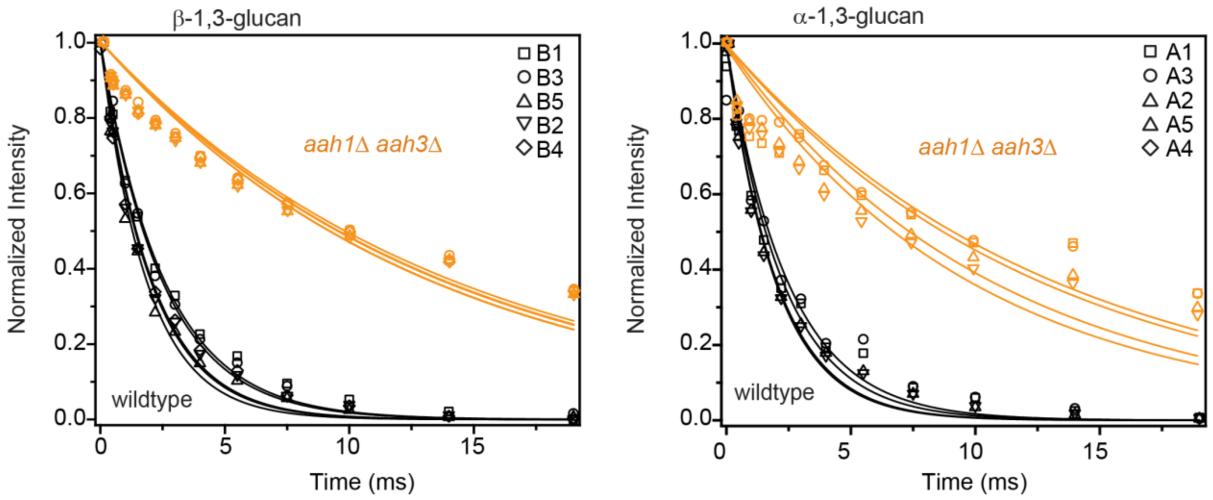


**Figure S5**. **$^1$H-T$_{1\rho}$ NMR relaxation curves with single-exponential fitting.** (**A**) $^1$H-T$_{1\rho}$ relaxation curves fit to single-exponential equations were plotted for different carbon sites in β-1,3-glucan (top) and α-1,3-glucan (bottom) as comparisons between wildtype cells (black) and *aah1*Δ *aah3*Δ mutant (yellow).

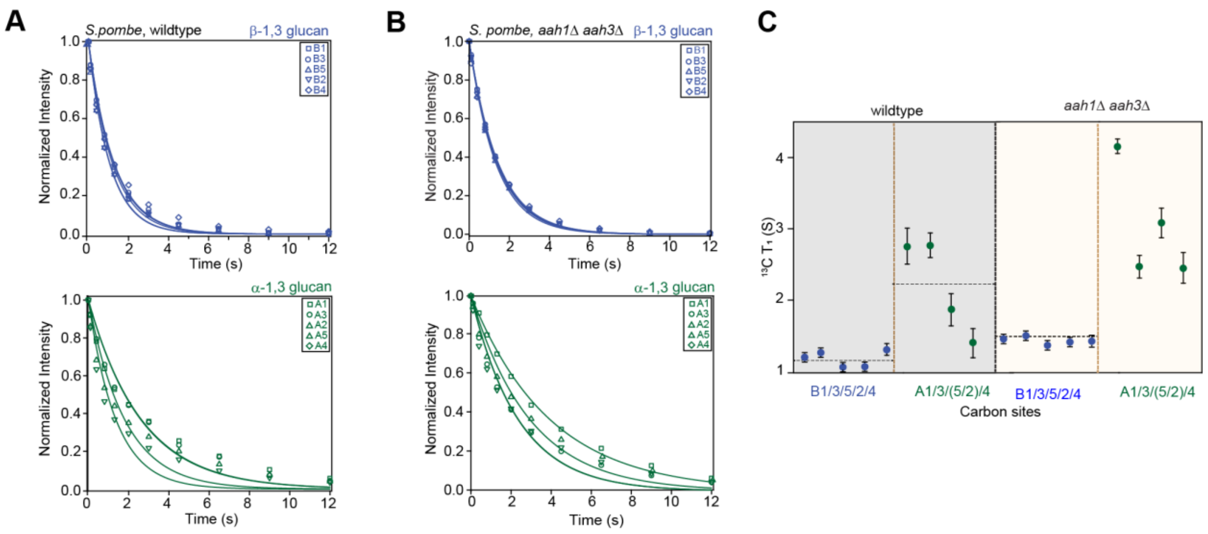


**Figure S6**. **$^{13}$C-T$_1$ NMR relaxation curves of polysaccharides with single-exponential fitting.** $^{13}$C-T$_1$ relaxation curves of β-1,3-glucans (blue) and α-1,3-glucans (green) are shown for (**A**) *wildtype* and (**B**) *aah1*Δ *aah3*Δ mutant. (**C**) $^{13}$C T$_1$ relaxation time constants for different carbon sites of β-1,3-glucan (blue) and α-1,3-glucan (green) in the wildtype (left; shaded in grey) and *aah1*Δ *aah3*Δ mutant (right; shaded in pale yellow). Symbols are used for assigning different carbons in the polysaccharides. The data are collected on 400 MHz (9.4 Tesla) spectrometer and the best fit is achieved using a single exponential equation.

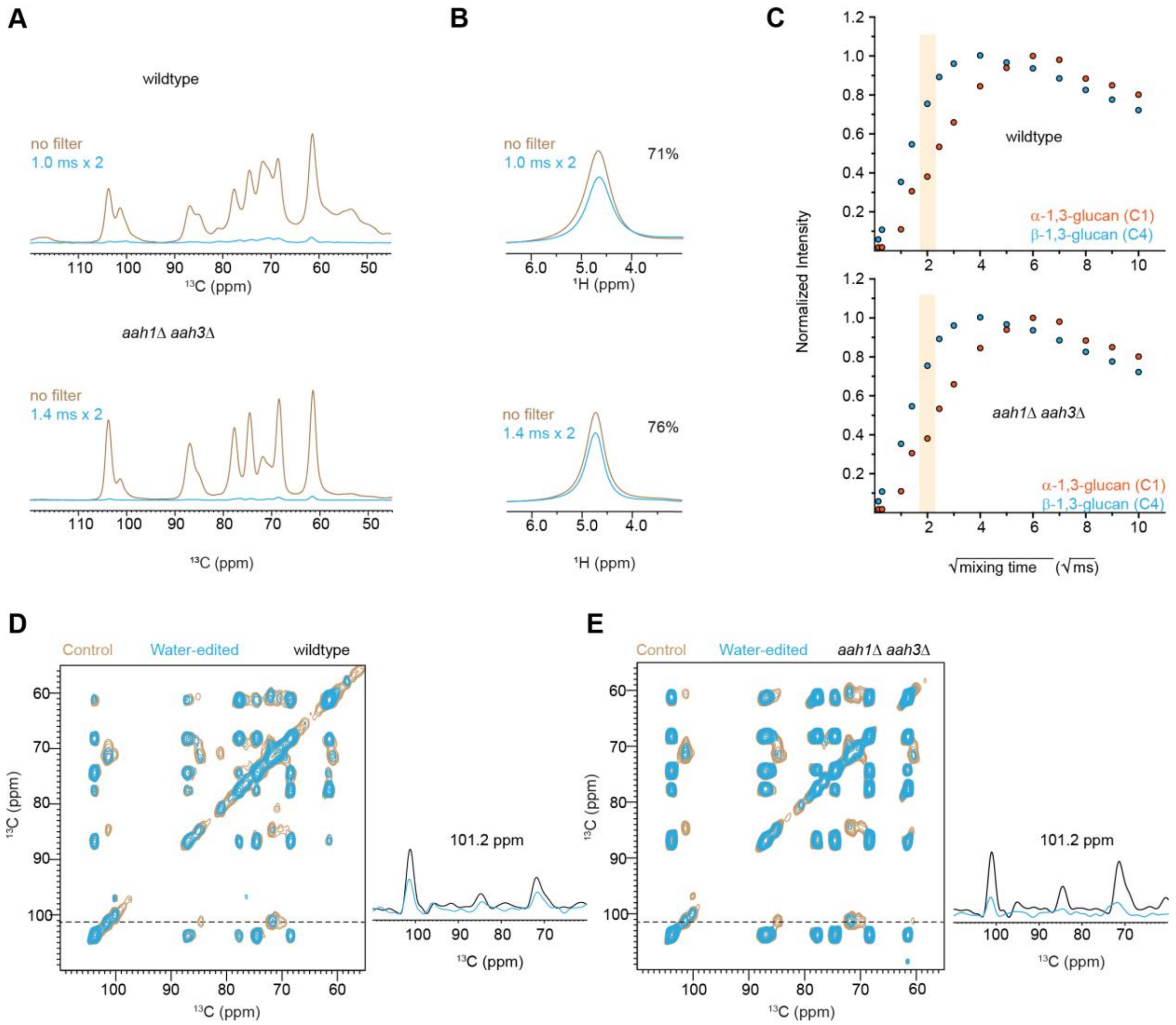


**Figure S7. Water-edited spectra of *S. pombe* to access polymer hydration.** (**A**) $^{1}$H-$T_2$ filtered (cyan) and control (almond) $^{13}$C spectra are shown for wildtype (top) and *aah1*Δ *aah3*Δ (bottom) *S. pombe* samples. No spin diffusion was applied. Approximately 95% of carbohydrate $^{13}$C signals were removed by the $^{1}$H-$T_2$ filter. (**B**) $^{1}$H-$T_2$ filtered (cyan) and control (almond) $^{1}$H NMR spectra, with 71% and 76% of water signal retained for wildtype and *aah1*Δ *aah3*Δ respectively after the $^{1}$H-$T_2$ filter. (**C**) Representative water-to-polysaccharide $^{1}$H spin diffusion buildup curves are shown for wildtype (top) and *aah1*Δ *aah3*Δ (bottom). Overlay of 2D water-edited (cyan) and control (orange) $^{13}$C-$^{13}$C correlation spectra of (**D**) wildtype sample and (**E**) *aah1*Δ *aah3*Δ mutant. Representative 1D slices extracted from the 2D $^{13}$C-$^{13}$C correlation spectra are shown for each sample. The control data are displayed as black, and the water-edited spectra are plotted in cyan. All spectra were measured on a 400 MHz spectrometer at 15 kHz MAS.

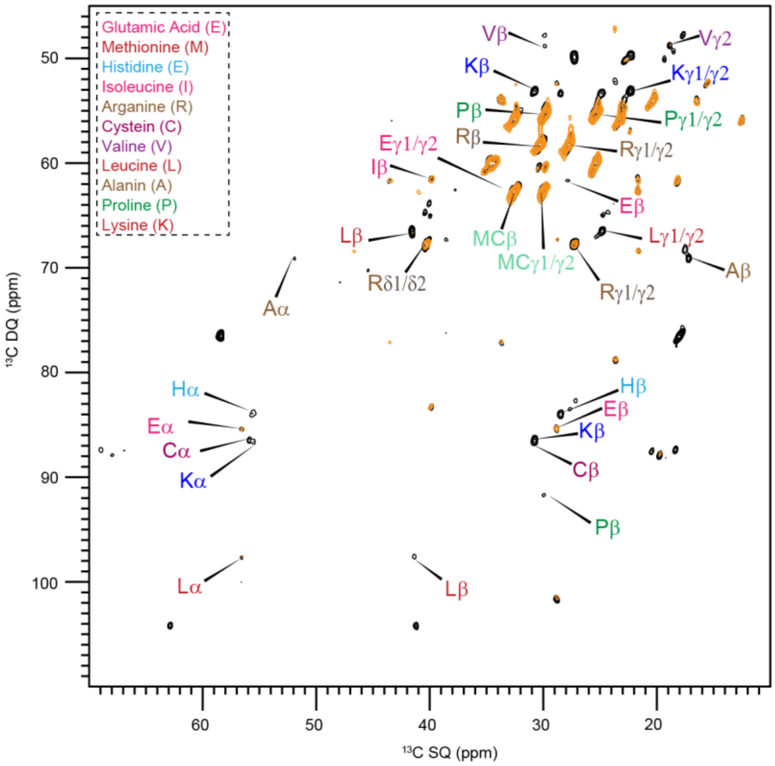


**Figure S8. Protein signals in *S. pombe***. The protein signals from the mobile phase analyzed from 2D $^{13}$C DP refocused J-INADEQUATE spectra. All the refocused INADEQUATE spectra were measured on 800 MHz spectrometer at 15 kHz MAS**.**

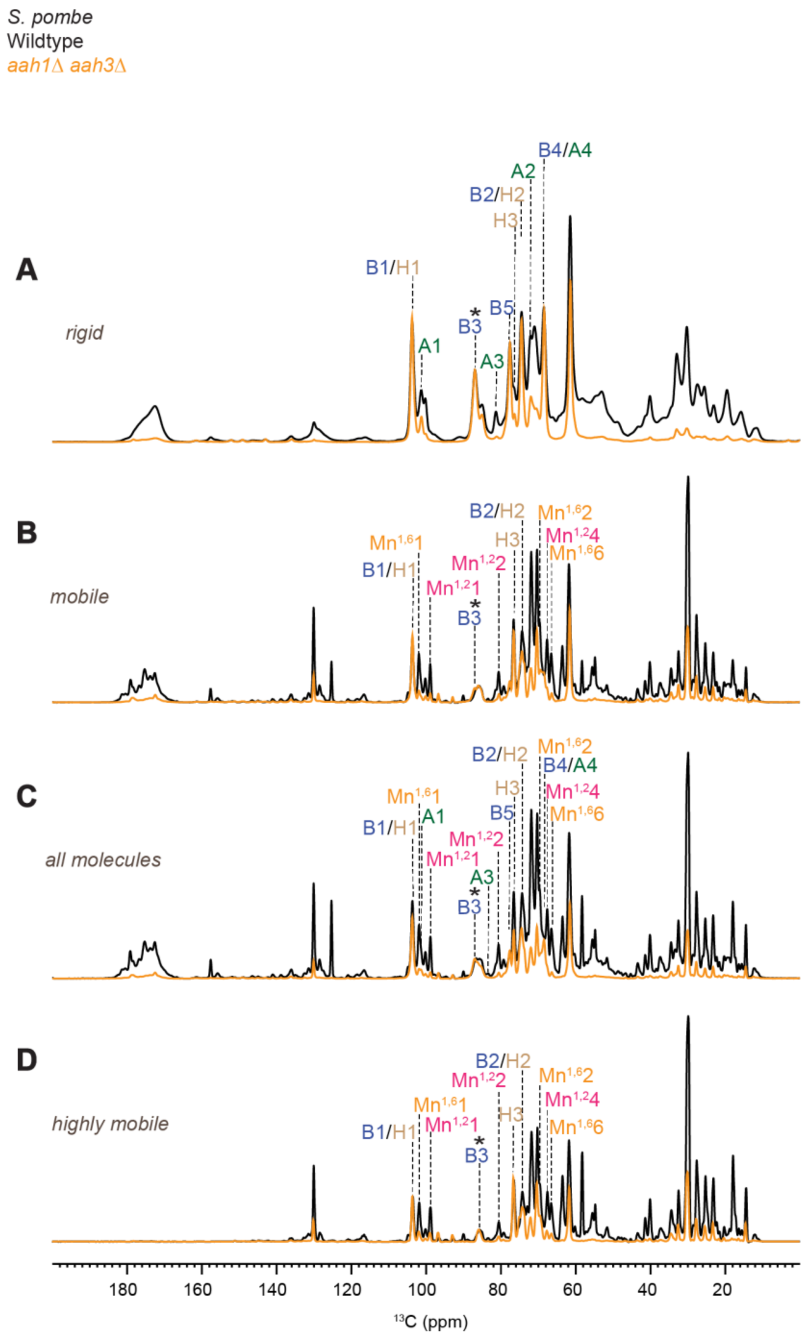


**Figure S9. Dynamical gradient of polysaccharides in *S. pombe* cell wall.** From top to bottom are four sets of 1D $^{13}$C spectra measured with (**A**) CP for selecting rigid polysaccharides. (**B**) DP spectra with a short recycle of 2s for selection of mobile components. (**C**) DP with long recycle delays (35s) for quantitative detection of all molecules. (**D**) Refocused INEPT experiments for probing the most dynamic molecules. The spectra for the wildtype of *S. pombe* and *aah1*Δ *aah3*Δ mutant is shown in black and orange color, respectively. Abbreviations: B, β-1,3-glucan; H, β-1,6-glucan; A, α-1,3-glucan; $Mn^{1,2}$, α-1-2-mannose; $Mn^{1,6}$, α-1,6-mannan.

**Table S1.** ***S. pombe*** **strains used in this study**

| Strain | Genotype | Source |
|---|---|---|
| **Figure 2** | | |
| KGY246 | *ade6-M210 leu1-32 ura4-D18* $h^-$ | Lab stock |
| KGY4728-2 | *aah1Δ::kanMX6 ade6-M210 leu1-32 ura4-D18* $h^-$ | Bioneer V3 |
| KGY3943-2 | *aah3Δ::kanMX6 ade6-M210 leu1-32 ura4-D18* $h^+$ | Bioneer V3 |
| KGY5335-2 | *aah1Δ::kanMX6 aah3Δ::kanMX6 ade6-M210 leu1-32 ura4-D18* $h^-$ | This study |
| **Fig. S4** | | |
| KGY16854 | *cwg1-1 ade6-M21X leu1-32 ura4-D18* $h^+$ | Ribas et al., 1991 |
| KGY5572-2 | *cwg1-1 aah3Δ::kanMX6 ade6-M210 leu1-32 ura4-D18* $h^-$ | This study |
| KGY17141 | *cps1-191 ade6-M210 leu1-32 ura4-D18* $h^+$ | Lab stock |
| KGY5574-2 | *cps1-191 aah3Δ::kanMX6 ade6-M210 leu1-32 ura4-D18* $h^+$ | This study |
| KGY2440 | *ags1-664 leu1-32* $h^-$ | Katayama et al., 1999 |
| KGY5522-2 | *ags1-664 aah3Δ::kanMX6 ade6-M210 leu1-32 ura4-D18* $h^+$ | This study |
| KGY1424-2 | *aah1Δ::kanMX6 pck2Δ::*$ura4^+$ *ade6-M210 leu1-32 ura4-D18* $h^+$ | This study |
| KGY1429-2 | *aah3Δ::*$ura4^+$ *pck2Δ::*$ura4^+$ *ade6-M210 leu1-32 ura4-D18* $h^-$ | This study |
| KGY1148-2 | *aah1Δ::kanMX6 pmk1Δ::*$ura4^+$ *ade6-M210 leu1-32 ura4-D18* $h^+$ | This study |
| KGY1088-2 | *aah3Δ::*$ura4^+$ *pmk1Δ::*$ura4^+$ *ade6-M210 leu1-32 ura4-D18* $h^-$ | This study |
| **Fig. S1** | | |
| KGY5153-2 | *aah2Δ::kanMX6 ade6-M210 leu1-32 ura4-D18* $h^-$ | This study |
| KGY3944-2 | *aah4Δ::kanMX6 ade6-M210 leu1-32 ura4-D18* $h^+$ | Bioneer V3 |
| KGY5275-2 | *aah1Δ::kanMX6 aah2Δ::kanMX6 ade6-M210 leu1-32 ura4-D18* $h^-$ | This study |
| KGY4939-2 | *aah1Δ::kanMX6 aah4Δ::kanMX6 ade6-M210 leu1-32 ura4-D18* $h^-$ | This study |
| KGY5277-2 | *aah2Δ::kanMX6 aah3Δ::kanMX6 ade6-M210 leu1-32 ura4-D18* $h^-$ | This study |
| KGY5279-2 | *aah2Δ::kanMX6 aah4Δ::kanMX6 ade6-M210 leu1-32 ura4-D18* $h^-$ | This study |
| KGY4941-2 | *aah3Δ::kanMX6 aah4Δ::kanMX6 ade6-M210 leu1-32 ura4-D18* $h^-$ | This study |
| KGY5337-2 | *aah1Δ::kanMX6 aah2Δ::kanMX6 aah3Δ::kanMX6 ade6-M210 leu1-32 ura4-D18* $h^-$ | This study |
| KGY5341-2 | *aah1Δ::kanMX6 aah2Δ::kanMX6 aah4Δ::kanMX6 ade6-M210 leu1-32 ura4-D18* $h^-$ | This study |
| KGY5339 | *aah1Δ::kanMX6 aah3Δ::kanMX6 aah4Δ::kanMX6 ade6-M210 leu1-32 ura4-D18* $h^-$ | This study |
| KGY5320-2 | *aah2Δ::kanMX6 aah3Δ::kanMX6 aah4Δ::kanMX6 ade6-M210 leu1-32 ura4-D18* $h^-$ | This study |
| **Fig. S2** | | |
| KGY896-2 | *sad1-mCherry:natMX6 ade6-M210 leu1-32 ura4-D18* $h^-$ | Lab stock |

**Table S2. Solid-state NMR experiments and parameters.** To be quantitative, direct pulse (DP) experiments with 35 s long recycling delay were used. cross-polarization (CP), most rigid molecules. With DP and a shorter recycling delay of 2 seconds, suppress the rigid molecules from the spectra, and with Insensitive Nuclei Enhanced by Polarization Transfer (INEPT) the most mobile molecules were selected. For 2D $^{13}C$-$^{13}C$ correlation experiments allowed to resolve rigid intramolecular peaks. 2D DQ-SQ, DP J-INADEQAUTE and CP INADEQUATE spectra were used to detect through-bond correlations. The experimental parameters include the $^{1}H$ Larmor frequency, total experiment time (t), recycle delay (d1), number of scans (NS), The number of points for the direct (td2) and indirect (td1) dimensions, the acquisition time of the direct dimension (aq2) and the evolution time of indirect dimension (aq1), spectral width (sw1 and sw2), mixing time ($t_m$), increment delay (IN_F) and T filter times. * Indicates the water-polysaccharide spin diffusion. The processing parameters include the window function and associated parameters.

| | Acquisition parameters | | | | | | | | | | | | | Processing parameters | |
|---|---|---|---|---|---|---|---|---|---|---|---|---|---|---|---|
| Experiment | $\omega_{0,1H}$ (MHz) | t (h) | d1 (s) | NS | td2 | td1 | aq2 (ms) | aq1 (ms) | sw2 (ppm) | sw1 (ppm) | $t_m$ (ms) | IN_F (µs) | T filters | Window function | Parameter |
| 1D $^{13}C$ CP | 800 | 0.5 | 2.0 | 1024 | 3200 | | 16.0 | | 496.8 | | 1.0 | | | GM | LB-10, GB0.05 |
| 1D $^{13}C$ DP | 800 | 0.2 | 2.0 | 256 | 3200 | | 16.0 | | 496.8 | | | | | GM | LB-10, GB0.05 |
| 1D $^{13}C$ DP | 800 | 2.5 | 35.0 | 256 | 3200 | | 16.0 | | 496.8 | | | | | GM | LB-10, GB0.05 |
| 1D $^{13}C$ INEPT | 800 | 1.5 | 3.5 | 1024 | 3200 | | 16.0 | | 496.8 | | | | | GM | LB-10, GB0.05 |
| 1D $^{13}C$ T1 | 400 | 3.5 | 2.0 | 512 | 2000 | | 16.0 | | 623.3 | | | | $T_1$($10^{-3}$-12 s) | GM | LB-10, GB0.05 |
| 1D $^{1}H$ T1ρ | 400 | 4.0 | 2.0 | 512 | 2000 | | 16.0 | | 623.3 | | | | SL ($10^{-3}$-19 ms) | GM | LB-10, GB0.05 |
| 2D $^{13}C$-$^{13}C$ CORD | 800 | 11.0 | 2.0 | 32 | 2800 | 600 | 14.0 | 7.5 | 496.8 | 198.7 | 53.0 | 25 | | QSINE | SSB 3.5 |
| 2D $^{13}C$-DP INADEQUATE | 800 | 6.0 | 2.0 | 16 | 2800 | 680 | 14.0 | 7.4 | 496.8 | 225.8 | | 22 | | | |
| 2D $^{13}C$-$^{13}C$ Water-edited | 400 | 8.0 | 2.0 | 64 | 2000 | 220 | 16.0 | 5.4 | 623.3 | 199.4 | $10^{-4}$ /50* | 50 | $T_2$ $10^{-4}$ ms | QSINE | SSB 3.5 |

**Table S3.** **$^{13}$C chemical shifts of biomolecules in *S. pombe* cell walls at ambient temperature.** Superscripts are used to denote different allomorphs. Not applicable (/). Unidentified (-). Branched (Br). Reducing end (O).

| Carbohydrate | | C1 | C2 | C3 | C4 | C5 | C6 | CO | $CH_3$ | N | Experiment | References |
|---|---|---|---|---|---|---|---|---|---|---|---|---|
| α-1,3-glucan | a | 101.1 | 72.2 | 84.9 | 68.5 | 69.1 | 61.2 | / | / | / | $^{13}$C-$^{13}$C CORD | Bhanja *et al*. 2014[1] |
| | b | 99.9 | 71.0 | 81.1 | 73.1 | 71.9 | 60.8 | / | / | / | $^{13}$C DP J-INADEQUATE | |
| | c | 101.9 | 68.4 | 81.4 | 69.5 | 71.8 | 60.9 | / | / | / | | |
| β-1,3-glucan | | 103.8 | 74.6 | 86.9 | 68.5 | 77.6 | 61.5 | / | / | / | $^{13}$C-$^{13}$C CORD | Shim *et al*. 2007[2] Fairweather *et al*. 2004[3] Saito *et al*. 1979[4] |
| β-1,3-glucan ($B^{Br}$) | | 103.2 | 73.9 | 85.6 | 69.1 | 76.1 | 69.2 | / | / | / | $^{13}$C DP J-INADEQUATE | Lowman et al. 2011[5] |
| β-1,6-glucan (H) | | 103.5 | 74.2 | 76.5 | 70.5 | 75.1 | 69.6 | / | / | / | | |
| $Mn^{1,2}$ | | 101.4 | 79.1 | 71.1 | 67.9 | 74.3 | 61.9 | / | / | / | $^{13}$C DP J-INADEQUATE | Latgé *et al*. 1994[6] Chakraborty *et al*. 2021[7] |
| Mn-$O^{1,2}$ | | 99.1 | 80.5 | 71.8 | 67.8 | 74.0 | 62.2 | / | / | / | | |
| $Mn^{1,6}$ | | 101 | 72.9 | 73.8 | 67.9 | 72.8 | 66.6 | / | / | / | | |
| Unknown (unk) | | 107.6 | 82.1 | 77.7 | - | - | - | / | | | | |

| Amino Acids | Cα | Cβ | Cγ/γ1 | Cγ2 | Cδ/δ1 | | Amino Acids | Cα | Cβ | Cγ/γ1 | Cγ2 | Cδ/δ1 | References |
|---|---|---|---|---|---|---|---|---|---|---|---|---|---|
| Glutamic Acid (E) | 55.2 | 27.2 | 33.8 | | | | Valine (V) | | 29.2 | 18.4 | | | Fritzsching *et al*. 2013[8] |
| Methionine (M) | | 32.0 | 29.5 | | | | Leucine (L) | 54.1 | 40.1 | 24.6 | | 22.4 | |
| Histidine (H) | | 27.9 | | | | | Alanine (A) | 51.3 | 16.6 | | | | |
| Arginine (R) | | 29.5 | 26.8 | | 39.5 | | Proline (P) | 61.1 | 29.1 | 24.8 | | | |
| Cysteine (C) | 55.0 | 30.5 | | | | | Lysine (K) | 55.1 | 30.9 | 21.5 | | | |
| Isoleucine (I) | | 36.1 | | 15.0 | | | | | | | | | |

**Table S4. $^1$H-T$_{1\rho}$ relaxation time constants with double-exponential fitting.** A double exponential equation was used to fit the $^1$H-T$_{1\rho}$ of β-1,3-glucan and α-1,3-glucan polysaccharides in *S. pombe* and the equation used is $I(t) = Ae^{-t/T_{1A}} + Be^{-t/T_{1B}}$, where $A + B = 1$. Error bars are standard deviations of the fit parameters.

| | | **wildtype** | | | | ***aah1*Δ *aah3*Δ mutant** | | | |
|---|---|---|---|---|---|---|---|---|---|
| | Cross peaks | A | $^1$H-T$_{1\rho}$ (ms) | B | $^1$H-T$_{1\rho}$ (ms) | A | $^1$H-T$_{1\rho}$ (ms) | B | $^1$H-T$_{1\rho}$ (ms) |
| β-1,3-glucan | B1 | 0.53 | 4.37±1.01 | 0.47 | 1.34±0.37 | 0.85 | 19.43 ± 0.84 | 0.15 | 0.76±0.17 |
| | B3 | 0.40 | 4.57±2.77 | 0.60 | 1.65±0.68 | 0.86 | 19.79 ± 1.16 | 0.14 | 0.87±0.27 |
| | B5 | 0.33 | 4.22±2.08 | 0.67 | 1.23±0.31 | 0.84 | 19.37 ± 1.22 | 0.16 | 0.86±0.25 |
| | B2 | 0.39 | 4.28±1.40 | 0.61 | 1.23±0.27 | 0.83 | 19.65 ± 1.22 | 0.17 | 1.04±0.27 |
| | B4 | 0.47 | 3.96±0.99 | 0.53 | 1.11±0.26 | 0.85 | 19.48 ± 0.88 | 0.15 | 0.77±0.18 |
| α-1,3-glucan | C1 | 0.45 | 4.89±1.23 | 0.55 | 1.12±0.26 | 0.78 | 22.79 ± 2.79 | 0.22 | 0.48±0.19 |
| | C3 | 0.47 | 5.07±1.80 | 0.53 | 1.20±0.42 | 0.83 | 19.99 ± 1.94 | 0.17 | 0.46±0.22 |
| | C2/C5 | 0.40 | 4.41±1.23 | 0.60 | 1.16±0.23 | 0.76 | 18.89 ± 1.73 | 0.24 | 0.91±0.24 |
| | C4 | 0.36 | 4.61±1.30 | 0.64 | 1.16±0.21 | 0.75 | 17.76 ± 1.73 | 0.25 | 0.88±0.23 |

**Table S5. $^{13}$C-$T_1$ and $^{1}$H-$T_{1\rho}$ relaxation time constants with single exponential fitting.** A single exponential equation was used to fit the $T_1$ data $I(t) = e^{-t/T_I}$. A single exponential equation was used to fit the $T_{1\rho}$ data: $I(t) = e^{-t/T_{I\rho}}$. Error bars are standard deviations of the fit parameters.

| Sample Type | Cross peaks | $T_1$ (s) | Cross peaks | $T_{1\rho}$ (ms) |
|---|---|---|---|---|
| Wildtype | B1 | 1.2±0.04 | B1 | 2.6±0.1 |
| | B3 | 1.3±0.04 | B3 | 2.5±0.2 |
| | B5 | 1.1±0.05 | B5 | 1.9±0.1 |
| | B2 | 1.1±0.05 | B2 | 2.0±0.1 |
| | B4 | 1.3±0.08 | B4 | 2.0±0.1 |
| | A1 | 2.6±0.20 | A1 | 2.3±0.1 |
| | A3 | 2.6±0.20 | A3 | 2.5±0.2 |
| | A2/A5 | 1.8±0.20 | A2/A5 | 2.0±0.1 |
| | A4 | 1.4±0.20 | A4 | 2.0±0.1 |
| *aah1*Δ *aah3*Δ mutant | B1 | 1.4±0.02 | B1 | 13.7±1.1 |
| | B3 | 1.5±0.01 | B3 | 14.1±1.1 |
| | B5 | 1.3±0.03 | B5 | 13.2±1.1 |
| | B2 | 1.4±0.03 | B2 | 13.2±1.1 |
| | B4 | 1.4±0.05 | B4 | 13.7±1.1 |
| | A1 | 3.9±0.10 | A1 | 12.6±1.7 |
| | A3 | 2.4±0.20 | A3 | 13.2±1.4 |
| | A2/A5 | 2.9±0.20 | A2/A5 | 10.7±1.1 |
| | A4 | 2.3±0.20 | A4 | 9.7±1.0 |
